# Combined effects of climate warming and pharmaceuticals on a tri-trophic freshwater food web

**DOI:** 10.1101/2023.03.31.535078

**Authors:** Claire Duchet, Kateřina Grabicová, Vojtech Kolar, Olga Lepšová, Helena Švecová, Andras Csercsa, Barbora Zdvihalová, Tomáš Randák, David S. Boukal

## Abstract

Multiple anthropogenic stressors influence the functioning of ponds and lakes, but their combined effects are often little understood. We ran two mesocosm experiments to evaluate the effects of warming (+4°C above ambient) and environmentally relevant concentrations of a mixture of commonly used pharmaceuticals, an emerging class of chemical contaminants, on tri-trophic food webs representative of pelagic communities in ponds and other small standing waters. We quantified the main and interactive effects of warming and pharmaceuticals on each trophic level and attributed them to the direct effects of both stressors and the indirect effects arising through biotic interactions. Warming and pharmaceuticals had stronger effects in the summer experiment, altering zooplankton community composition and causing delayed or accelerated emergence of top insect predators. In summer, both stressors and top predators reduced filter-feeding zooplankton biomass, while warming and pharmaceuticals had opposing effects on phytoplankton. In the winter experiment, the effects were much weaker and primarily limited to a positive effect of warming on phytoplankton biomass. Overall, we show that pharmaceuticals can exacerbate the effects of climate warming in freshwater ecosystems, especially during the warm season. Our results demonstrate the utility of community-level studies across different seasons for the risk assessment of multiple emerging stressors in freshwater ecosystems.

## Introduction

Freshwater ecosystems face severe impacts from multiple stressors, including habitat loss, chemical pollution, pathogens, invasive species, and climate change (IPBES, 2019). Among these stressors, climate warming and chemical pollution pose major threats to freshwater biodiversity and ecosystem functioning (Vörösmarty et al., 2010). Climate change is predicted to increase average global annual temperatures by up to +4.4°C by 2100, with stronger warming at high latitudes and during cold seasons (Masson-Delmotte et al., 2021). Freshwater ecosystems, dominated by ectotherms, are highly susceptible to warming (Parmesan and Yohe, 2003; Woodward et al., 2010) that can lead to reduced body sizes, altered species interactions, and shifts in species distribution ranges and phenology, with cascading impacts on ecosystem functioning (Boukal et al., 2019; Post, 2013).

Among chemicals, pharmaceuticals represent a prominent class of emerging pollutants (Fekadu et al., 2019). Their widespread use and inadequate removal in wastewater treatment plants contribute to global contamination of aquatic systems (Wilkinson et al., 2022). At environmentally relevant concentrations (ng.L^-1^ to low µg.L^-1^), pharmaceuticals can disrupt the behavior (Brodin et al., 2014; Saaristo et al., 2019) and reproduction of aquatic ectotherms (Fursdon et al., 2019), affecting individual fitness and triggering indirect effects like altered trophic interactions (Fursdon et al., 2019; Bláha et al., 2019). Although pharmaceuticals can bioaccumulate in benthic macroinvertebrates (Grabicova et al., 2015), leading to exposure of higher trophic levels (Lagesson et al., 2016) like fish (Brodin et al., 2014; Du et al., 2014; Grabicova et al., 2017), they are usually diluted through the trophic level and rarely biomagnified (Grabicova et al., 2015; Xie et al., 2017).

The effects of pharmaceuticals on aquatic systems have primarily been studied using single-species laboratory bioassays, often at higher concentrations than those found in the environment (Richards et al., 2004). Furthermore, pharmaceuticals typically occur as complex mixtures in the environment (Grabicova et al., 2015; Loos et al., 2013). Understanding their ecotoxicology under environmentally realistic conditions and quantitative consequences on the biota therefore requires new data (Backhaus, 2014). Moreover, the combined effects of pharmaceuticals and warming on freshwater biota and ecosystem functioning remain largely unknown (Heye et al., 2019) despite extensive research on their individual effects. While the dominant stressor alone often explains the main impacts of multiple stressors on a freshwater ecosystem (the so-called dominance null model; Morris et al., 2022), interactions between stressors can result in synergistic, antagonistic, or additive effects, making predictions challenging (Spaak et al., 2017).

Warming can modulate the effects of pollutants on freshwater ecosystems both directly and indirectly. While many pollutants become more toxic at higher temperatures, warming may also accelerate their degradation, thereby reducing the exposure of biota to them (Noyes et al., 2009; Op de Beeck et al., 2017). Population-level responses to the combined stressors can vary dramatically due to differences in environmental sensitivity (Venâncio et al., 2016), tolerance of individuals (Baert et al., 2016), ecological trade-offs and local adaptation patterns (Bennett and Lenski, 2007; Kneitel and Chase, 2004). Additionally, the fate of species embedded in food webs depends on direct and indirect changes in intraspecific and interspecific interactions (Post, 2013; Relyea, 2009), including trait- and density-mediated indirect effects such as top-down and bottom-up effects (Brogan and Relyea, 2015; Guedes et al., 2016; Rodrigues et al., 2018). Finally, persistent pollutants can alter community structure and affect or enhance various aspects of ecosystem functioning (Duchet et al., 2018; Sánchez-Bayo, 2021).

To address the knowledge gap on the combined effects of pharmaceuticals and warming on freshwater communities, we conducted a full-factorial experiment using outdoor mesocosms. By simulating realistic conditions, we aimed to provide insights into the ecological consequences of these stressors in a changing environment. We included key functional groups representative of the communities in pond and shallow lake ecosystems and ran the experiment during both summer and winter to account for seasonal turnover in the communities. Our pelagic communities included three main trophic levels and we therefore expected (1) a strong negative effect of warming on zooplankton and none or a positive effect on the top predators (Hansson et al., 2013), leading to an increase in phytoplankton biomass (Belden et al., 2007). We also expected that (2) warming will have stronger negative effects during summer and milder, more positive effects during winter, especially on phytoplankton and zooplankton growth.

## 2. Material and Methods

### 2.1. Mesocosms

The mesocosm facility at the University of South Bohemia (České Budějovice, Czech Republic; 48°58’36.6” N 14°26’41.8” E, 381 m a.s.l.) comprises 32 circular outdoor mesocosms (120 cm inner diameter, 1.13 m3 water volume each) with a custom-built heating system (Hennlich®, Schädling, Austria; Fig. 1). The 28 mesocosms used in this study were filled 6 weeks before the experiment with tap water filtered through a granulated activated carbon filter, enriched with 100 g of common alder leaves (*Alnus glutinosa*), and 8 g of fish food (Pondstick granules, Apetit®) to add nutrients. Water in each mesocosm was gently mixed with an air stone, except from 12 January to 3 February 2021, when mixing was paused to allow natural freezing in the unheated mesocosms (Fig. S1).

**Figure 1.**
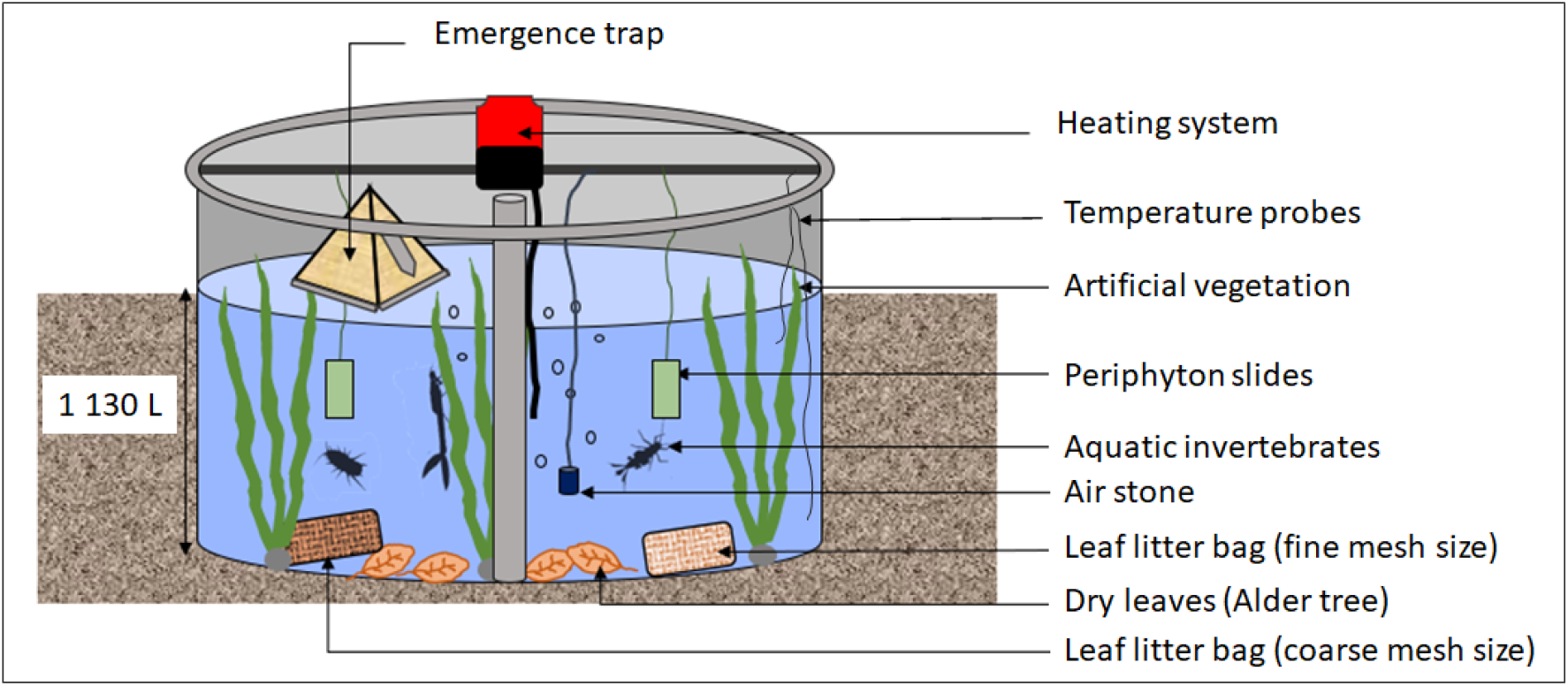
Schematic view of the mesocosm set-up

The experimental community included common macroinvertebrate and plankton taxa representing different trophic levels and feeding guilds (predators, filter feeders, grazers, detritivores, and primary producers; Table S2). Phytoplankton and zooplankton from a nearby fishpond were added to the mesocosms one month before the experiment, after collecting water samples for xenobiotic analysis (see detailed methods in S3, Tables S3.2 and S3.3). The plankton inoculum was mixed in a barrel and then added to the mesocosms (2.5-litre aliquot each). Macroinvertebrates from nearby fishponds, ditches, and sand pits were then sequentially introduced into the mesocosms after two weeks: benthic detritivores (*Asellus aquaticus* - ca. 100 per mesocosm), grazers (*Cloeon* cf. *dipterum* mayfly larvae - ca. 200 per mesocosm), omnivorous scrapers (*Planorbarius corneus* pond snails - 9 per mesocosm), omnivorous piercers (*Sigara falleni* water boatman - 50 per mesocosm), water column predators (*Notonecta glauca* backswimmers - 9 individuals per mesocosm, *Aeshna* or *Anax* dragonfly larvae - 6 per mesocosm; hereafter Aeshna and Anax), benthic predators (libellulid dragonfly larvae - 3 per mesocosm), and phytophilous predators occupying artificial vegetation and mesocosm walls (damselfly larvae - ca. 50 per mesocosm). The predators were added to the mesocosms one week before the start.

Each mesocosm received three artificial submerged macrophytes made of four strips of green plastic mesh (5×100 cm) attached to a granite stone to provide additional microhabitat. Two polypropylene strips (6 cm wide, 50 cm long) hanging vertically in each mesocosm were used to incubate pre-cultured periphyton from one of the fishponds where the macroinvertebrates were collected. One coarse (20×20 cm; 20 mm mesh) and one fine (20×20 cm; 0.5 mm mesh) litter bag containing 9.5 g of dried alder (*Alnus glutinosa*) leaves were added to each mesocosm to distinguish the contribution of detritivores and microbial decomposers to the decomposition process (Fig. 1).

### 2.2. Experimental design

Seven mesocosms were assigned to each of the four treatments: (1) unheated without pharmaceuticals, (2) unheated with pharmaceuticals, (3) heated without pharmaceuticals, and (4) heated with pharmaceuticals. Each mesocosm received the same treatment in both experiments. Heating was maintained at +4°C above ambient throughout each experiment (mean ± SD differences in daily mean temperatures between treatments: winter experiment, 4.03 ± 0.04°C; summer experiment, 4.06 ± 0.06°C; see S1), consistent with the SSP5-8.5 projection model for 2081–2100 (Masson-Delmotte et al., 2021).

The pharmaceutical mixture included 15 compounds from four major drug categories (cardiovascular: telmisartan, valsartan, metoprolol, atenolol; psychoactive: carbamazepine, lamotrigine, venlafaxine, citalopram, tramadol; antihistaminic: cetirizine, fexofenadine; antibiotic: sulfamethoxazole, trimethoprim, clarithromycin, clindamycin; Table S3.1) at concentrations commonly detected in surface waters in the Czech Republic (Fedorova et al., 2022; Švecová et al., 2021). The pharmaceutical mixture was administered as a single pulse exposure (total concentration: 2890 ng.L^-1^; see S3 for more details).

We conducted two experiments: the ‘winter experiment’ from 24 September 2020 to 12 March 2021 and the ‘summer experiment’ from 2 June to 2 August 2021 (Figures S4.1 and S4.2). Day 0 marked the start of each experiment before adding the pharmaceuticals and initiating warming. The initial three samplings for all parameters were performed on Day 0, followed by Day 2 and Day 7. Subsequent samples were collected monthly in winter and fortnightly in summer.

### 2.3. Sampling

This section covers the methods for assessing the temporal changes in the pelagic environment and biota including predatory insects feeding on zooplankton. Complete community data collected at the end of each experiment will be analyzed elsewhere. Table S4.1 provides a summary of all response variables collected during the experiments.

In both experiments, we measured pharmaceutical concentrations, environmental parameters (dissolved O_2_, conductivity, pH, turbidity, water temperature, and chlorophyll-a concentration) using a portable multiparameter water quality meter (YSI 556 MPS probe; YSI®, USA). On each sampling day, one water sample (6 mL) was collected in each mesocosm, filtered through a 0.2-µm syringe filter (regenerated cellulose) and stored at - 20°C until the pharmaceutical compound concentrations were analysed with an in-line solid-phase extraction liquid chromatography with tandem mass spectrometry (in-line SPE/LC-MS/MS, triple quadrupole mass spectrometer Quantiva, Thermo Fisher Scientific, USA; see S3 for details).

Dissolved organic carbon (DOC), nitrogen, and phosphorus concentrations were measured at Day 0 and at the end of each experiment. For technical reasons, total nitrogen (TN) and total phosphorus (TP) were measured during the winter experiment, while dissolved nitrogen (DN) and soluble reactive phosphorus (SRP) were measured during the summer experiment. At the end of each experiment, phytoplankton samples were collected from the surface water layer (ca. 10 cm) in 500 ml PET bottles and fixed with Lugol solution immediately after sampling before identification and counting in the lab (see S4.1 for details).

During both experiments, we repeatedly sampled zooplankton with a tube sampler (1.2 m length, 7 cm inner diameter) equipped with a one-way bottom valve. On each sampling date, three samples were taken from different locations within each mesocosm to minimize plankton patchiness effects (Stephenson et al., 1984). The resulting bulk sample (approximately 9 L depending on water level, ranging between 7.5–9.6 L in summer and 8.4– 10.5 L in winter) was collected in a 20 L bucket. From this, one litre was filtered through an 80-μm mesh plankton net. Retained organisms (zooplankton and some other pelagic invertebrates) were preserved in 4% formaldehyde in 120 mL plastic vials. Counting and identification were done under a stereomicroscope (Olympus SZX2-ILLT stereomicroscope; Olympus, Tokyo, Japan) to the lowest feasible taxonomic level using identification keys (Amoros, 1984; Johannsen, 1937).

During the summer experiment, we also monitored aquatic insect emergence with pyramid-shaped floating emergence traps (35 × 35 cm base dimensions) with a 250 mL collection bottle of soapy water (Cadmus et al., 2016). One trap was installed in each mesocosm on Day 0. Emerging insects were collected every 2–3 days and preserved in 70% ethanol (mayflies, chironomids) or frozen (odonates) for later identification and analysis of pharmaceutical content.

Biomass of each taxon was estimated by multiplying total abundance (for aquatic insects) or density (zooplankton) by the average body size based on literature data and our measurements (Table S4.2).

### 2.4. Statistical analyses

We employed multiple methods (detailed in S4.2) to assess the effects of the pharmaceutical mixture and warming on the aquatic community. In brief, redundancy analysis (RDA) was used to examine temporal changes in environmental parameters and chlorophyll-a concentrations. Principal component analyses (PCAs) were used to visualize differences in algal groups and taxa between treatments. Principal response curve (PRC) was used to analyse changes in zooplankton community structure. We then ran generalized linear models (GLMs) to assess treatment effects on nutrients, predator emergence and survival and generalized linear mixed models (GLMMs) to assess treatment effects on *Daphnia* and copepod biomasses. Model selection approach with the corrected Akaike information criterion was used to identify the most parsimonious model for each univariate response. Finally, we used confirmatory path analysis (piecewise SEM) to understand the direct and indirect effects of the pharmaceutical mixture and warming on the community. Hypothetical pathways were based on *a priori* knowledge to build a complete model of cause-effect relationships. Fisher’s C statistics were used to assess the overall fit of each SEM model. Multivariate analyses were performed using CANOCO 5 (ter Braak and Smilauer, 2012), and univariate analyses were done in R version 4.1.2 (R Core Team, 2018).

## 3. Results

### 3.1. Pharmaceutical concentrations

During the summer experiment, total pharmaceutical concentration gradually decreased over time to about 25% in heated mesocosms and 30% in non-heated mesocosms (Fig. 2A). Meanwhile, metabolite concentrations increased during the first month and remained stable thereafter (Fig. 2C). In the winter experiment, pharmaceuticals were degraded over time (Fig. 2B). Contrarily, metabolite concentrations in the winter experiment increased significantly during the first two months, reaching double the summer concentrations, and remained stable afterward (Fig. 2D). Pharmaceuticals and their metabolites were below detection limits in samples from non-treated mesocosms in both experiments.

**Figure 2.**
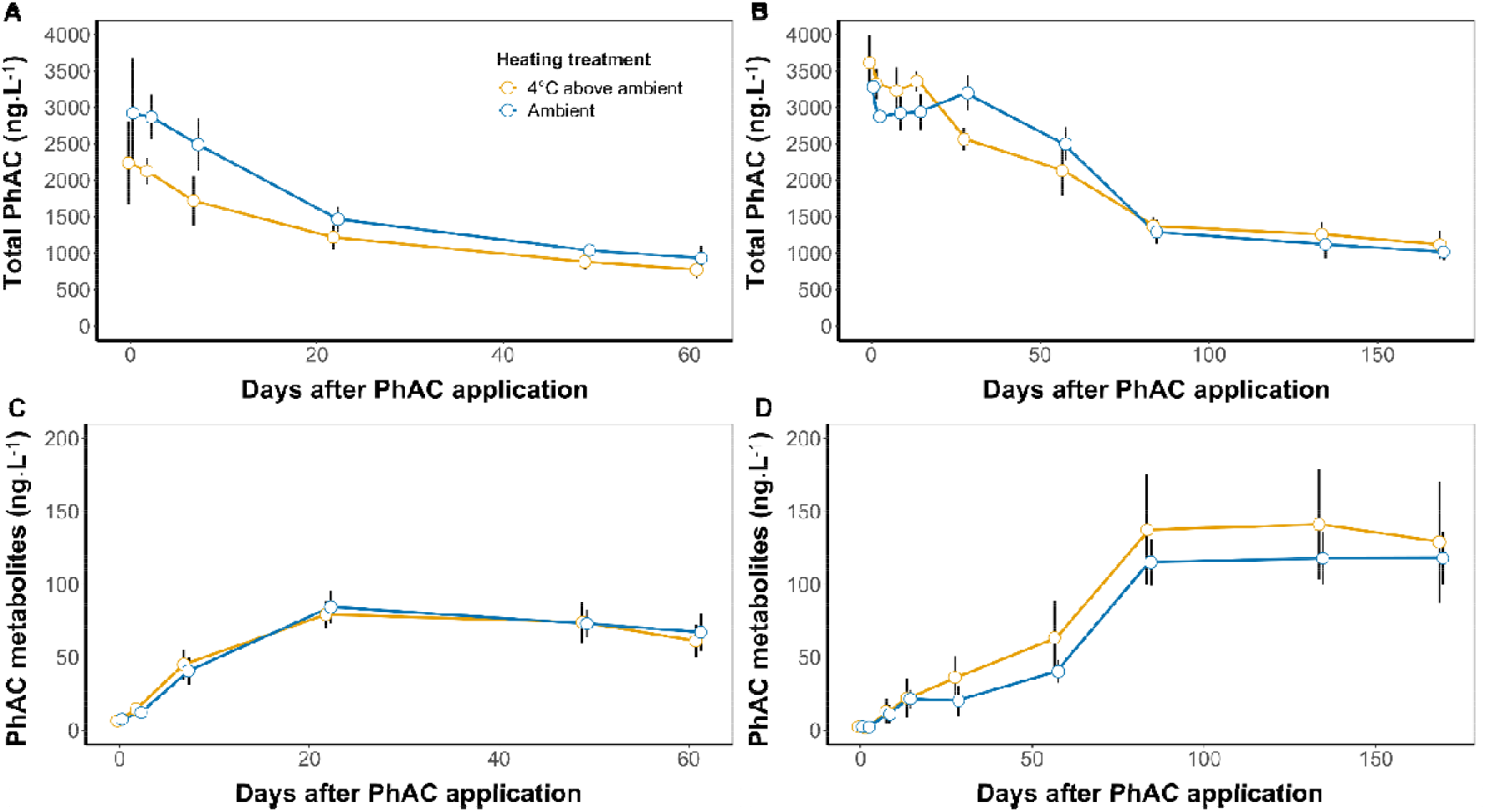
Total concentrations of pharmaceutical (PhAC) main compounds (A, C) and their metabolites (B, D) in the winter (A, B) and summer (C, D) experiments in heated (orange) and non-heated (blue) mesocosms treated with pharmaceuticals. Symbols with error bars = treatment-specific mean values ± SD (*n* = 7); mean values in the same treatment connected by colour lines.

### 3.2. Environmental conditions

Environmental conditions significantly differed between treatments in both experiments (summer RDA: pseudo-F = 4.9, p = 0.002, explained variation = 7.4%; winter RDA: pseudo-F = 53.7, p = 0.001, explained variation = 43.1%). In the summer experiment, non-heated mesocosms had higher water levels, higher turbidity, lower conductivity, and lower chlorophyll-a concentrations (mainly those without pharmaceuticals) compared to heated mesocosms (Fig. 3A, Table S5.1). In the winter experiment, treatments did not differ in chlorophyll-a concentration, dissolved oxygen, turbidity, and pH, but non-heated mesocosms had higher water levels and lower conductivity compared to the heated ones (Fig. 3B, Table S5.2).

**Figure 3.**
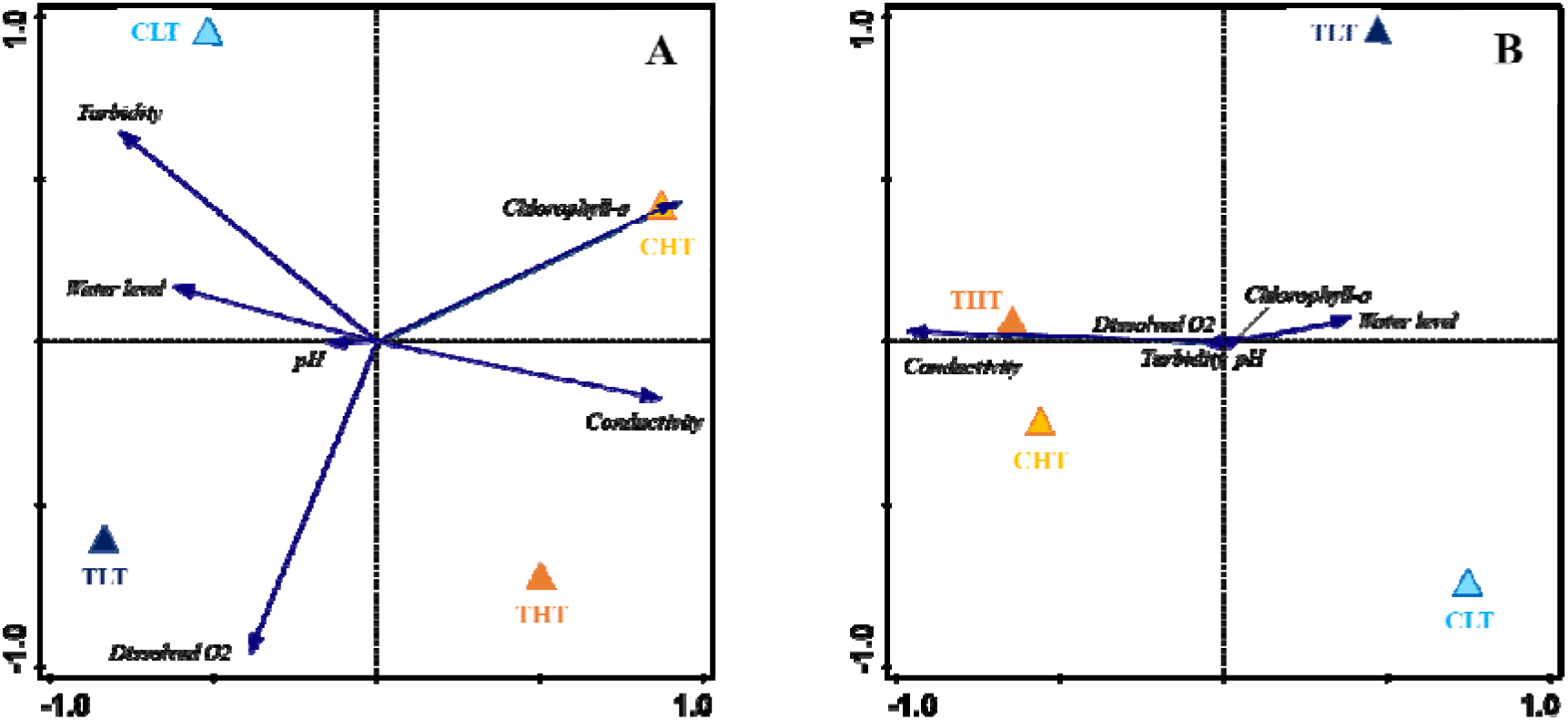
Ordination diagram (RDA) showing differences between the treatments in environmental parameters and chlorophyll-a concentration during the summer (A) and winter (B) experiment. Treatments: CLT = non-heated, without pharmaceuticals; CHT = heated, without pharmaceuticals; TLT = non-heated, with pharmaceuticals; THT = heated, with pharmaceuticals.

### 3.3. Effect of the stressors on nutrients and phytoplankton

The most parsimonious models showed disparate effects of the stressors on nutrients in the summer and winter experiments. In summer, heated mesocosms had lower final concentrations of DOC, DN, and SRP compared to non-heated ones. The added pharmaceutical mixture had a small negative effect on DOC and a significant negative effect on DN and SRP (Fig. S6.1A, C, and E). In winter, heated mesocosms also had lower final concentrations of DOC, TN, and TP compared to non-heated ones, but no effects of pharmaceuticals on nutrients were detected (Fig. S6.1B, D, and F). No synergistic effects of the two stressors on nutrients were detected in both experiments (Tables S6.1 and S6.2).

There were some differences in the composition of major algal groups between treatments (PCA, Fig. S6.2), although nonsignificant (RDA, Table S6.3). In summer, Cyanobacteria dominated in the heated mesocosms, while Chrysophyta were prevalent in the non-heated mesocosms (Fig. S6.2A-B). In winter, Chrysophyta were more prevalent in the non-heated controls, while Streptophyta were more prevalent in the non-heated, treated treatment (Fig. S6.2C). Warming appeared to be the main driver of taxonomic composition in the winter experiment (PCA F2, Fig. S6.2D). Notably, chlorophytes (*Oocystis* spp. and *Scenedesmus* sp.) were more prevalent in the heated mesocosms, while bacillariophytes (*Synedra* sp., *Gomphonema* spp., *Navicula* sp. and *Ulnaria* sp.) and chrysophytes were relatively more abundant in the non-heated mesocosms (Fig. S6.2D).

### 3.4. Effect of the stressors on zooplankton community dynamics

The dynamics and composition of the zooplankton community and its responses to the stressors differed markedly between the summer and winter experiments (Fig. 4). In summer, zooplankton biomass showed significant differences between treatments (PRC: F = 0.7, p = 0.003; Fig. 4A). These differences were mainly driven by warming, with noticeable effects on ostracods and *Daphnia* (Fig. 4A). Ostracod biomass increased in heated mesocosms, while *Daphnia* biomass decreased compared to non-heated mesocosms (Figs 4A and S7.1b, Table S7.3). However, copepod biomass tended to increase in all treatments, with a higher increase in the treated mesocosms (Fig. S7.2a, Table S7.3). In winter, there were no significant differences in zooplankton biomass and composition between treatments (F = 0.3, p = 0.41; Fig. 4B, Figs S7.1c and S7.2b; Table S7.4).

**Figure 4.**
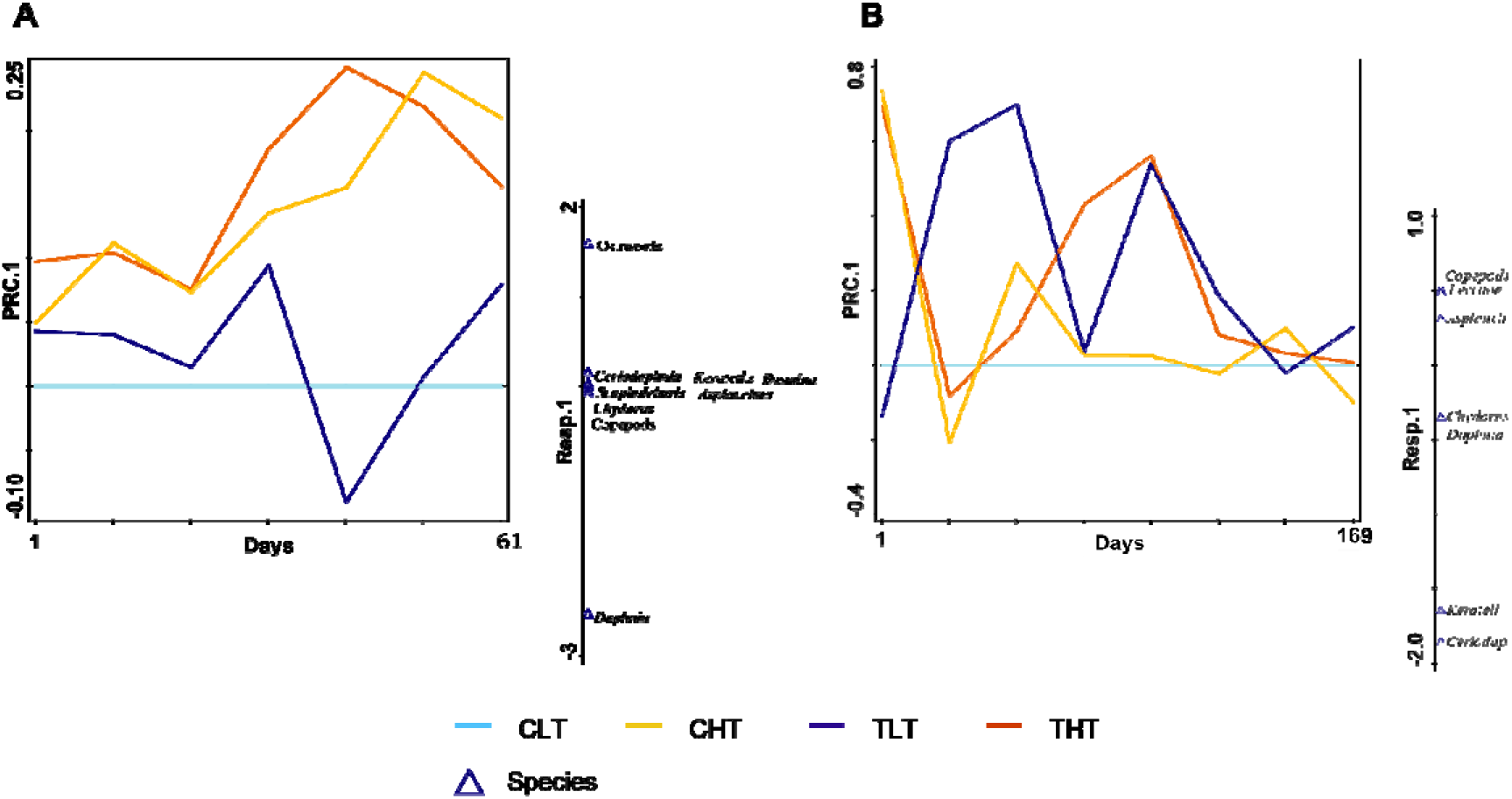
Principal response curves (PRC) with species weights (vertical bars) indicating treatment-specific effects on zooplankton during the summer (A) and winter (B) experiment. Days = days after exposure. Treatments: CLT = non-heated, without pharmaceuticals (light blue); CHT = heated, without pharmaceuticals (yellow); TLT = non-heated, with pharmaceuticals (dark blue); THT = heated, with pharmaceuticals (orange). Days = days after exposure.

### 3.5. Effect of the stressors on predatory insect survival and emergence

The effects of both stressors on emergence and survival of predatory insects differed between taxa and experiments (Fig. 5; Tables S8.1 and S8.2). In summer, both warming and pharmaceuticals increased the emergence of anisopteran (*Aeshna*) larvae, with an additive effect on the predictor scale (Fig. 5A). However, warming had a negative effect on damselfly emergence (Fig. 5B). In winter, the survival of anisopteran (*Anax*) larvae was similar in all treatments, while damselfly larvae showed better survival when exposed to pharmaceuticals only or heating only, resulting in similar survival in treatments without both stressors and with both stressors (Fig. 5C). *Notonecta* survival was reduced when exposed to warming in winter, but pharmaceuticals had no effect on their survival (Fig. 5D).

**Figure 5.**
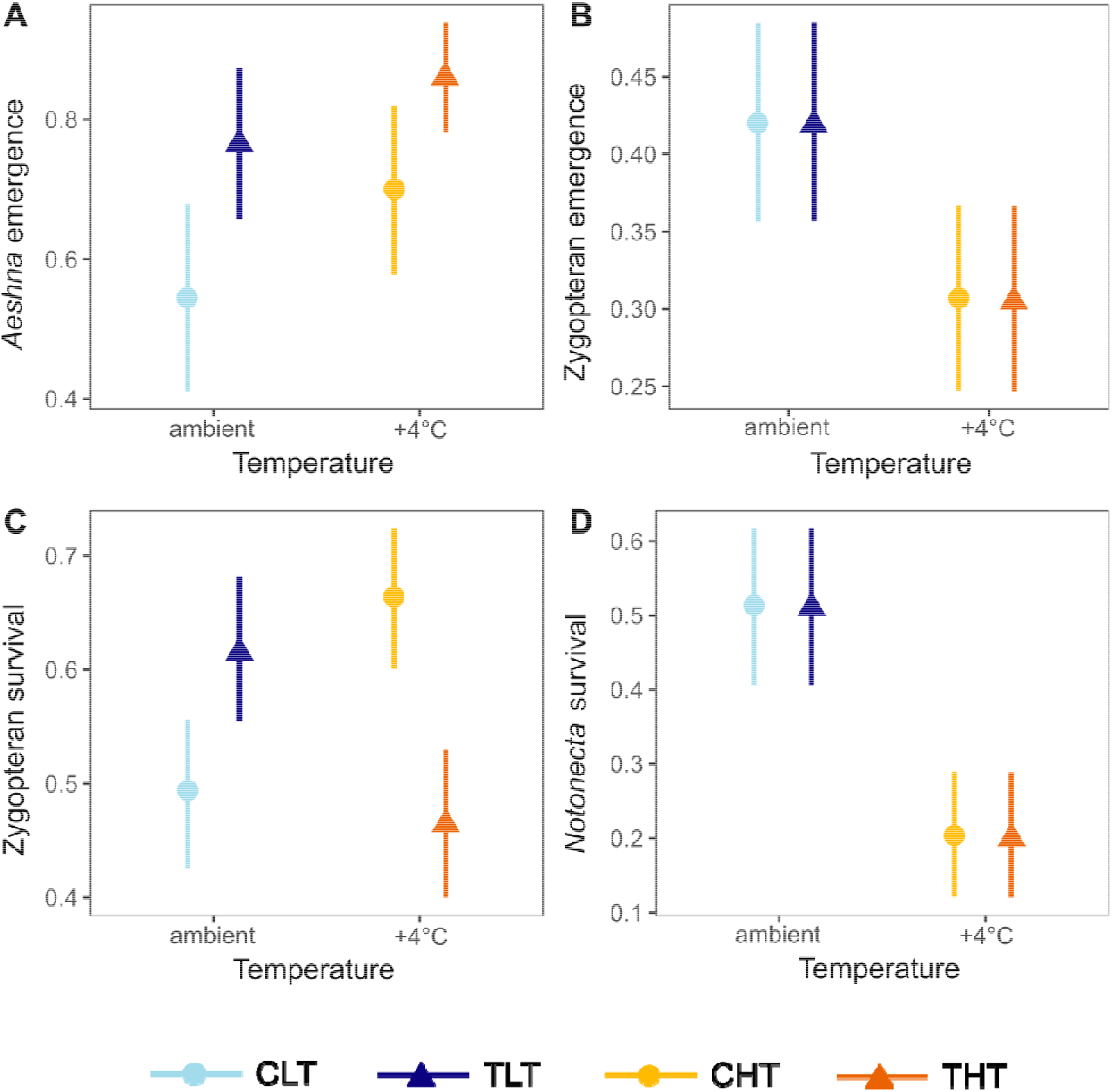
Estimates of treatment-specific probability of emergence of insect predators in the summer experiment (A: anisopteran larvae (*Aeshna*), B: zygopteran larvae) and probability of survival of insect predators in the winter experiment (C: zygopteran larvae, D: *Notonecta* backswimmers). Treatments: CLT = non-heated, without pharmaceuticals; CHT = heated, without pharmaceuticals; TLT = non-heated, with pharmaceuticals; THT = heated, with pharmaceuticals. Model estimates are based on the most parsimonious models and include 95% confidence intervals based on fixed effects.

### 3.6. Effect of the stressors on the pelagic food web

The effects of the stressors on the pelagic food web differed between summer and winter experiments (Fig. 6 and Tables S9.1 and S9.2). In summer, pharmaceuticals negatively affected the phytoplankton (standardised path coefficient < −0.01; Fig. 6A and Table S9.1), while warming had a positive direct effect (standardised path coefficient = +0.08). Both stressors had a direct and negative impact on filter-feeding cladocerans (standardised path coefficients < −0.01 for pharmaceuticals and = −0.14 for warming). However, the relationship between primary producers and filter feeders was not significant (standardised path coefficient = −0.09), contradicting the trophic cascade hypothesis. The effects of stressors on predators were positive; it was significant for warming (direct effect, standardised path coefficients = +0.02) but not for pharmaceuticals (direct effect, standardised path coefficients < 0.01). Finally, predators directly caused a decrease in filter feeder biomass (standardised path coefficient = −0.21; Fig. 6A).

**Figure 6.**
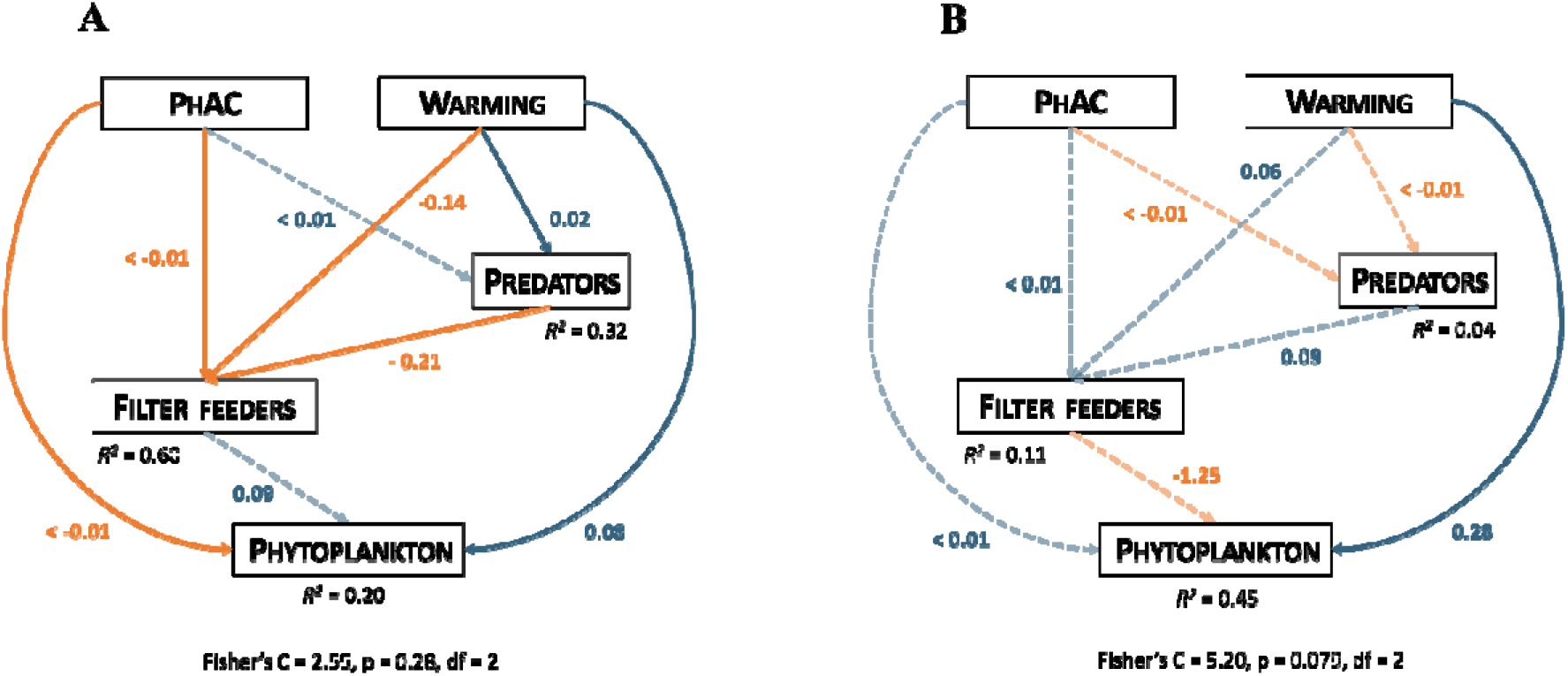
Path diagram of the selected structural equation model based on average data collected during the summer experiment, and last sampling data during the winter experiment. PhAC = pharmaceuticals. Orange, blue, and dashed arrows respectively indicate significantly (*p* < 0.05) negative, significantly positive, and non-significant (*p >* 0.05) relationships between variables. Proportions of variation explained by the model for each response variable given by *R*^2^ values. See Tables S9.1 and S9.2 for details.

In contrast, in winter, the pharmaceutical mixture had no significant direct or indirect effect on any component of the simplified pelagic food web. Only phytoplankton biomass was directly and positively affected by warming during winter (standardized path coefficient = 0.28; Fig. 6B and Table S9.2).

## 4. Discussion

We conducted the first study on the combined effects of a complex mixture of pharmaceuticals and warming on freshwater communities. While previous studies focused on simplified food webs (Wijewardene et al., 2021), we examined the consequences of these stressors on three trophic levels, including phytoplankton, zooplankton, and aquatic insects as top predators.

As expected, we observed only limited effects of the stressors on the community in the winter experiment, whereas in summer, warming acted as the stronger stressor in line with the dominance null model (Morris et al., 2022), and its effects amplified the effects of pharmaceuticals on the freshwater community. Overall, both stressors had a negative effect on zooplankton (mainly composed by filter feeders in our experiment) through direct and indirect effects, promoting phytoplankton development. Our results demonstrate that the effects of environmentally relevant pharmaceutical mixtures should be considered together with warming when studying possible future responses of complex freshwater communities to environmental stressors.

### 4.1. Effects of warming on the freshwater food web

Warming had the stronger impact on the freshwater food web in our experiments, especially during summer. During the summer experiment, the daily mean temperature in the heated mesocosms reached up to 29.5°C (compared to 25.5°C in the non-heated mesocosms) and remained above 25°C for almost three weeks in July. Such temperatures are comparable to the optimum growth temperatures of the larvae of many odonates (Suhling et al., 2015) and green algae (25–35 °C range; Bouterfas et al., 2002) but well above the reported optima for *Daphnia* (16–22 °C range; Bruijning et al., 2018).

Warming may indirectly impact community dynamics through increased evaporation, potentially leading to eutrophication and shifts in stoichiometry (Ptacnik et al., 2010). In our summer experiment, water levels decreased faster in the heated mesocosms due to higher evaporation. However, our experimental designs prevented hypoxic and anoxic conditions, and no significant stoichiometry shift in available nutrients was observed at the end of the experiment.

Warming can also favour fast-growing copiotrophs (Mooij et al., 2008; Ren et al., 2017; Zhang et al., 2019) and toxin-producing cyanobacteria (Huisman et al., 2018), intensifying resource competition with phytoplankton (Joint et al., 2002). Moreover, it can reduce phytoplankton diversity, leading to shifts in community composition (Urrutia-Cordero et al., 2017). In our summer experiment, phytoplankton biomass (indicated by chlorophyll-a concentrations) increased, with cyanobacteria becoming more prevalent in the heated mesocosms towards the end of the summer experiment, while non-heated mesocosms remained dominated by green algae. These findings align with Hansson et al. (2020), who showed that a +4°C temperature increase led to a shift in the phytoplankton community, favouring cyanobacteria.

Zooplankton grazing can modulate the effects of warming on phytoplankton (Velthuis et al., 2017). For example, Yvon-Durocher et al. (2011) found that a +4°C temperature increase in an outdoor mesocosm experiment led to a decrease in phytoplankton biomass, favouring smaller organisms within the phytoplankton assemblage, while zooplankton biomass remained unaffected unlike in our experiment.

In tri-trophic systems, the effects of warming should vary predictably between trophic levels (Hansson et al., 2013). Top predators, by consuming zooplankton and reducing grazing pressure on phytoplankton, often play a crucial role in modulating the impacts of climate warming on the community (Hansson et al., 2013; Šorf et al., 2015; Velthuis et al., 2017). Consequently, warming is expected to have positive effects on top predators and phytoplankton but negative effects on zooplankton (Hansson et al., 2013 but see Yvon-Durocher et al., 2011). For example, He et al. (2018) observed interactive effects of warming (+3°C above ambient) and predation (by the fish *Aristichthys nobilis*) in an outdoor mesocosm experiment, leading to higher phytoplankton biomass and lower biomasses of *Daphnia* and total zooplankton under the combined stressors.

Results from our summer experiment support and further extend these findings. We observed a shift in the zooplankton community composition under warming, with ostracods dominating in response to higher temperatures. Predator survival and emergence also increased under warming. Moreover, predation rate of odonate larvae increases with warmer temperatures (e.g. Sentis et al., 2015), resulting in accelerated development and earlier emergence (Suhling et al., 2015). We conclude that these mechanisms caused higher predation pressure on the zooplankton, which reduced its grazing pressure and led to higher phytoplankton biomass.

### 4.2. Effects of pharmaceuticals on the freshwater food web

We observed no significant effects of pharmaceuticals in the winter experiment. However, the pharmaceutical mixture alone had a negative effect on nutrients, phytoplankton biomass and *Daphnia* biomass during summer.

Pharmaceutical mixtures can significantly impact microbial and algal communities and ectotherms from zooplankton to fish (Brodin et al., 2014). However, the negative effects reported in some studies were based on higher concentrations than those used in our study, potentially overestimating environmental consequences. For instance, a mixture of 10 pharmaceuticals was toxic to algae, daphniids, and zebrafish at concentrations up to 15,000 times higher than environmental levels (Watanabe et al., 2016). High concentrations of a pharmaceutical mixture (analgesics, antidepressants and antibiotics; >60 mg.L^-1^ for each compound) in outdoor microcosms caused fish mortality and reduced zooplankton and phytoplankton diversity (Richards et al., 2004). Conversely, David et al. (2020) found no strong effects on a set of biomarkers and population dynamics of three-spined sticklebacks (*Gasterosteus aculeatus*) at environmentally relevant concentrations of five pharmaceuticals (antiepileptics, analgesics, and antihypertensives).

In previous *in-situ* studies, antibiotic mixtures induced significant structural and functional changes in microbial attached communities (Proia et al., 2013), while a mixture of pharmaceuticals decreased algal biomass and photosynthesis (Rosi-Marshall et al., 2013). The decline in phytoplankton biomass exposed to pharmaceuticals in our summer experiment supports these findings.

### 4.3. Interactions between warming and pharmaceuticals

While warming had stronger effects in our experiments, the combined impact of the pharmaceutical mixture and warming was not predictable from individual stressor effects, particularly in the summer experiment. The combination of pharmaceuticals and warming significantly reduced phytoplankton biomass, and its control by filter-feeding zooplankton directly and indirectly via increased insect predator predation. This led to a change in phytoplankton composition, as expected in warming conditions (Kratina et al., 2012; Lewandowska et al., 2014). The qualitative effects of pharmaceuticals on top predators and zooplankton resembled those of warming in a tri-trophic system (Hansson et al., 2013), but their impact on phytoplankton differed.

These findings highlight the importance of community-level studies to assess stressor effects (Attrill and Depledge, 1997; Brooks and Crowe, 2019; Smith et al., 2009) and avoid underestimating their impacts (Jentsch et al., 2011). Relying solely on individual response variables for ecosystem function can be misleading in identifying underlying drivers of ecosystem changes (Bracken and Stachowicz, 2006; Hooper and Vitousek, 1998)

The interactions between temperature and contaminants present challenges for ecological risk assessment, particularly with global warming (Zhang et al., 2019). While the effects of warming on contaminant toxicity are well-studied, the impacts on shallow freshwater ecosystems can be complex, influenced by factors like contaminant type, community composition, and trophic interactions. For instance, heat waves combined with pesticide exposures can result in antagonistic or synergistic effects depending on the pesticides involved (Polazzo et al., 2022). Additionally, systems near their tolerance limit for one stressor may exhibit a lower threshold to withstand subsequent stress events (Hewitt et al., 2016).

Finally, the outcomes of our summer and winter experiments highlight the challenge of differentiating stress timing from seasonal species turnover. Although the communities had similar functional groups, differences in species composition, although subtle, made it challenging to disentangle the effects of seasonality, stressors, and community composition. However, we consider the taxonomic differences to be less significant than the overall seasonal stage of the community, with winter communities being dormant or exhibiting low metabolic activity compared to the highly active summer communities (see Kratina et al., 2012).

Brooks and Crowe (2019) suggested considering temporal variation in both the cause and effect to enhance understanding and management of multiple stressors. With increasing unpredictability in the future, accounting for the intensity and duration of stress events becomes crucial as they play a significant role in ecosystems’ responses (Easterling et al., 2000; Jentsch et al., 2007; Walter et al., 2013). Although our study did not directly assess seasonal variation, previous research indicates that climate-driven multiple stressors may vary seasonally (Godbold and Solan, 2013; Kratina et al., 2012). Additionally, climate change may alter terrestrial runoff patterns (Easterling et al., 2000; Masson-Delmotte et al., 2021), affecting local stressor arrival in aquatic systems (Adams, 2005; Schiedek et al., 2007). Further research is therefore required to explore the consequences of these changes.

## 5. Conclusions and perspective

Our findings demonstrate that low concentrations of pharmaceuticals alone had minimal impact, but when combined with warming, they significantly altered the pelagic food web, especially during summer when temperatures were close to or above the upper thermal limits for many species. Additionally, we observed distinct effects of warming and pharmaceuticals between seasons.

Our study highlights that the responses of freshwater communities to environmental stressors are complex and involve both direct effects on individual species and indirect effects through trophic interactions. These complexities are vital to consider when predicting future changes in biodiversity and ecosystem functioning (Hansson et al., 2013; Polazzo et al., 2022). The lack of community-level studies limits our understanding of ecosystem-level responses, especially when studying multiple interacting stressors. Although distinguishing between the effects of multiple stressors and understanding emergent phenomena is challenging, future research should consider larger spatio-temporal scales and a more comprehensive understanding of food web interactions. This will help us better predict the impacts of warming and emerging contaminants like pharmaceuticals and aid decision-making to mitigate their effects on freshwater ecosystems.

## 6. Acknowledgements

This study was supported by the Czech Science Foundation (project number GA20-16111S) and by the Ministry of Education, Youth and Sports of the Czech Republic (project CENAKVA, LM2023038). We thank Vladimíra Dekanová, Kateřina Bláhová, Bruno Carreira, Samuel Dijoux, Marius Wieczek and Fanny Verfaillie for assistance during the mesocosm experiment.

