## Supplementary material for "Combined effects of climate warming and pharmaceuticals on a tri-trophic freshwater food web"

**S1: Water temperature profiles in both experiments**

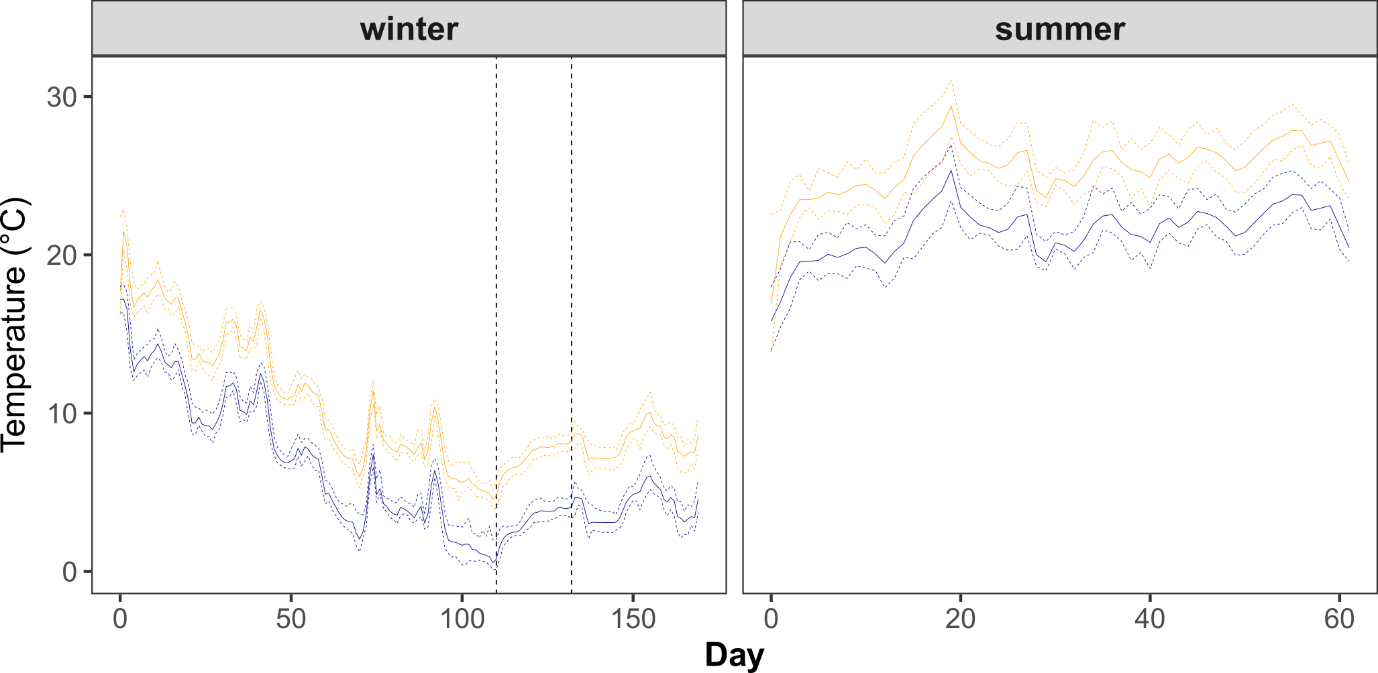

**Figure S1.** Temperature profiles (daily mean temperatures in bold line, minimum and maximum daily temperatures in dash lines) in the mesocosms during the winter and the summer experiments. Blue lines: non-heated mesocosm; yellow lines: heated mesocosms. Vertical dashed lines: start and end of the period when the water mixing was stopped.

**S2: Origin of the collected animals**

**Table S2.** **Macroinvertebrate taxa added to the mesocosms.** Feeding guild classification based on Schmidt-Kloiber and Hering (2015); quantity given per each mesocosm and represents individuals unless stated otherwise. Rare non-target taxa are omitted.

| **Taxon** | **Class** | **Feeding guild** | **Quantity** | **Site information** | **GPS coordinates** |
| --- | --- | --- | --- | --- | --- |
| *Lymnaea stagnalis* | Gastropoda | grazer / shredder | 8 | sandpit ponds  (Cep) | 48.9177011N, 14.8829131E |
| *Planorbarius corneus* | Gastropoda | grazer / shredder | 9 | artificial channel  (Zlata Stoka) | 49.0901736N, 14.7097089E |
| *Cloeon dipterum* | Insecta | grazer / scraper – gatherer / collector | 200 | sandpit ponds  (Cep) | 48.9177011N, 14.8829131E |
| *Sigara* sp. | Insecta | predator / collector | 50 | fishpond (Černiš) | 49.0032561N, 14.4339564E |
| Zygoptera | Insecta | Predator | 50 | sandpit ponds (Cep) and fishpond (Horni Machovec) | 48.9177011N, 14.8829131E  49.0038886N, 14.3696117E |
| *Aeshna* sp. (summer) | Insecta | Predator | 6 | small forest ponds (Řidka Blana) | 49.0829619N, 14.3914447E |
| *Anax* sp. (winter) | Insecta | Predator | 6 | sandpit ponds  (Cep) | 48.9177011N, 14.8829131E |
| *Sympetrum* sp. | Insecta | predator | 4 | sandpit ponds  (Cep) | 48.9177011N, 14.8829131E |
| *Notonecta* sp. | Insecta | predator | 9 | sandpit ponds  (Cep) | 48.9177011N, 14.8829131E |
| Zooplankton | Crustacea | filter feeder (cladocerans, rotifers) | 1L of inoculum | fishpond (Černiš) | 49.0032561N, 14.4339564E |
| *Asellus aquaticus* | Crustacea | grazer / scraper – gatherer / collector | 100 | artificial channel (Zlata Stoka) | 49.0901736N, 14.7097089E |

**S3: Chemical analyses**

The pharmaceutical mixture was administered in the mesocosms as a single pulse exposure of 20 ng.L^-1^ for citalopram and fexofenadine, 50 ng.L^-1^ for atenolol and clindamycin, 100 ng.L^-1^ for clarithromycin, sulfamethoxazole, trimethoprim and venlafaxine, 150 ng.L^-1^ for valsartan, 200 ng.L^-1^ for cetirizine, 250 ng.L^-1^ for carbamazepine and metoprolol and 500 ng.L^-1^ for lamotrigine, telmisartan and tramadol (total concentration: 2890 ng.L^-1^).

Water samples collected for the analysis of the pharmaceutical concentrations were stored at -20⁰C before analysis. Analytical methods followed (Grabicová et al., 2020; Koba et al., 2018; Lindberg et al., 2014) Acetonitrile (LC-MS grade; Merck), and ultra-pure water (Milli-Q® Direct 8, Merck, Germany) were used as mobile phases, acidified (by 0.1% HPLC/MS grade formic acid, Merck).

Twenty-six pharmaceuticals and their metabolites were used for preparing stock solution and as analytical standards (see Table S3.1; for detailed information on analytes and corresponding internal standards (see Grabicová et al., 2020; Koba et al., 2018). After thawing of filtered water samples at room temperature, internal standards were added, and the samples were analysed using in-line solid-phase extraction liquid chromatography with tandem mass spectrometry (in-line SPE/LC-MS/MS, triple quadrupole mass spectrometer Quantiva, Thermo Fisher Scientific, USA). Hypersil Gold aQ column (20 x 2.1 mm, 12 µm particles, Thermo Fisher Scientific) was used for analytes trapping, and Hypersil Gold aQ column (50 x 2.1 mm, 5 µm particles, Thermo Fisher Scientific) was used for chromatographic separation. For more details of the method, see (Grabicová et al., 2020; Koba et al., 2018; Lindberg et al., 2014). The limits of quantification (LOQs) and half-lives for parent compounds are reported in Table S3.1.

**Table S3.1: Compounds used for the pharmaceutical stock solution and for the analysis.** Main compounds given in bold and metabolites in normal characters, LC-MS/MS Mass Transitions, and limits of quantification (LOQs, ng/L, present as min-max values).

| **Analyte** | **Producer** | **Parent ion** | **Quan ion** | **LOQ min** | **LOQ max** |
| --- | --- | --- | --- | --- | --- |
| **Atenolol** | **Sigma Aldrich** | **267.17** | **190.0863** | **1.2** | **16** |
| **Carbamazepine (CBZ)** | **Sigma Aldrich** | **237.10** | **194.0967** | **0.25** | **11** |
| Oxcarbazepine | CDN Isotopes | 247.17 | 204.1590 | 0.43 | 33 |
| 10,11-transdihydro CBZ | TRC | 271.11 | 254.0813 | 0.84 | 26 |
| 10,11-dihydroCBZ | Fluka | 239.12 | 194.0965 | 0.44 | 11 |
| CBZ-10,11-epoxide | TRC | 253.10 | 210.0913 | 0.2 | 12 |
| **Cetirizine** | **Sigma Aldrich** | **389.16** | **201.0467** | **0.24** | **15** |
| **Citalopram** | **AK Scientific** | **325.17** | **109.0452** | **0.41** | **8.1** |
| N-desmethylcitalopram | Sigma Aldrich | 311.16 | 109.0450 | 0.42 | 8.6 |
| **Clarithromycin** | **Sigma Aldrich** | **748.50** | **158.1178** | **5** | **15** |
| **Clindamycin** | **Sigma Aldrich** | **425.19** | **126.1281** | **0.55** | **7.3** |
| Clindamycin sulfoxide | TRC | 441.18 | 377.1835 | 0.4 | 6.4 |
| **Fexofenadine** | **Sigma Aldrich** | **502.30** | **466.2742** | **0.38** | **13** |
| **Lamotrigine** |  |  |  | **4** | **32** |
| **Metoprolol** | **Sigma Aldrich** | **268.19** | **116.1072** | **0.59** | **11** |
| Metoprolol acid | TRC | 268.15 | 145.0648 | 0.26 | 19 |
| **Sulfamethoxazole (SMX)** | **Riedel-de Haen** | **254.06** | **156.0113** | **0.39** | **11** |
| N1-acetylSMX | CIL | 260.08 | 162.0316 | 0.9 | 34 |
| N4-acetylSMX | Sigma Aldrich | 296.07 | 198.0221 | 0.8 | 12 |
| **Telmisartan** | **AK Scientific** | **515.20** | **497.2335** | **1.6** | **52** |
| **Tramadol** | **Sigma Aldrich** | **264.20** | **58.0660** | **3.7** | **14** |
| Tramadol_IS, D3 | Lipomed | 267.21 | 58.0659 | 0.49 | 95 |
| **Trimethoprim** |  |  |  | **0.6** | **11** |
| **Valsartan** | **Sigma Aldrich** | **436.23** | **291.1490** | **0.45** | **7.1** |
| **Venlafaxine** | **AK Scientific** | **278.21** | **58.0657** | **0.39** | **8.3** |
| O-desmethylvenlafaxine | Sigma Aldrich | 264.20 | 246.1853 | 0.55 | 13 |

**Table S3.2.** **Xenobiotic concentrations in the sampling sites and the minimum and maximum LOQ values (ng.L^-1^).** LOQ_1_ = LOQ values for Horni Machovec, Cep and Zlata stoka sampling sites; LOQ_2_ = LOQ values for Černiš site. Pharmaceuticals used for the experiment appear in bold characters in the table.

|  | **Horni Machovec** | **Cep** | **Zlata Stoka** | **Černiš** | **LOQ_1_ min** | **LOQ_1_ max** | **LOQ_2_ min** | **LOQ_2_ max** |
| --- | --- | --- | --- | --- | --- | --- | --- | --- |
| *Alfuzosin* | *<2.6* | *<1.8* | *<2.3* | *<5.1* | *1.8* | *2.6* | *5.1* | *5.6* |
| *Alprazolam* | *<7.8* | *<7.4* | *<8.1* | *<9.7* | *7.4* | *8.1* | *9.7* | *11* |
| *Amitryptyline* | *<2.5* | *<2.4* | *<2.7* | *<2* | *2.4* | *2.7* | *2* | *2.3* |
| ***Atenolol*** | ***<7*** | ***<8.8*** | ***<9.2*** | ***<7.7*** | ***7*** | ***9.2*** | ***7.7*** | ***8*** |
| *Atorvastatin* | *<1.8* | *<1.7* | *<1.9* | *<8.2* | *1.7* | *1.9* | *8.2* | *9.3* |
| *Azithromycin* | *<1.7* | *<1.3* | *<1.5* | *<42* | *1.3* | *1.7* | *42* | *47* |
| *Bezafibrate* | *<2.3* | *<2.2* | *<2.4* | *<8.8* | *2.2* | *2.4* | *8.8* | *9.9* |
| *Biperiden* | *<2.6* | *<1.9* | *<2.1* | *<2.5* | *1.9* | *2.6* | *2.5* | *2.8* |
| *Bisoprolol* | *<2.2* | *<1.6* | *<2.4* | *<1.7* | *1.6* | *2.4* | *1.7* | *1.9* |
| *Caffeine* | *38* | *100* | *110* | *20* | *21* | *32* | *5.6* | *6.6* |
| ***Carbamazepine*** | ***<9.8*** | ***<9.3*** | ***<10*** | ***<9.7*** | ***9.3*** | ***10*** | ***9.7*** | ***11*** |
| ***Dihydro CBZ*** | ***<1.7*** | ***<1.6*** | ***<1.8*** | ***<9.8*** | ***1.6*** | ***1.8*** | ***9.8*** | ***11*** |
| ***Epoxy CBZ*** | ***<1.4*** | ***<1.3*** | ***<1.5*** | ***<10*** | ***1.3*** | ***1.5*** | ***10*** | ***11*** |
| ***trans-dihydro-***  ***dihydroxy CBZ*** | ***38*** | ***<10*** | ***<11*** | ***<9.5*** | ***10*** | ***11*** | ***9.5*** | ***11*** |
| ***Cetirizine*** | ***<2.1*** | ***<2*** | ***<2.2*** | ***<8.4*** | ***2*** | ***2.2*** | ***8.4*** | ***9.4*** |
| *Cilazapril* | *<2.2* | *<2.1* | *<2.3* | *<6.9* | *2.1* | *2.3* | *6.9* | *7.8* |
| ***Citalopram*** | ***<2.4*** | ***<2.1*** | ***<2.4*** | ***<9.3*** | ***2.1*** | ***2.4*** | ***9.3*** | ***10*** |
| ***N-Desmethylcitalopram*** | ***<2.9*** | ***<2.1*** | ***<3*** | ***<9.6*** | ***2.1*** | ***3*** | ***9.6*** | ***11*** |
| ***Clarithromycin*** | ***<16*** | ***<12*** | ***<14*** | ***<9.1*** | ***12*** | ***16*** | ***9.1*** | ***9.8*** |
| *Clemastine* | *<9.9* | *<8.4* | *<11* | *<2.4* | *8.4* | *11* | *2.4* | *2.7* |
| ***Clindamycin*** | ***<2*** | ***<2.1*** | ***6.8*** | ***<6*** | ***2*** | ***2.3*** | ***6*** | ***6.7*** |
| ***Clindamycin_sulfoxide*** | ***<1.3*** | ***<1.4*** | ***3*** | ***<5.8*** | ***1.3*** | ***1.5*** | ***5.8*** | ***6.5*** |
| *Clomipramine* | *<1.9* | *<2.3* | *<2.7* | *<1.8* | *1.9* | *2.7* | *1.8* | *2.1* |
| *Clonazepam* | *8.5* | *7.8* | *5.1* | *<3.1* | *2.2* | *2.6* | *3.1* | *3.6* |
| *Codeine* | *<5.5* | *<5.3* | *<5.8* | *<9.5* | *5.3* | *5.8* | *9.5* | *9.8* |
| *Diclofenac* | *<5.7* | *<6.1* | *<6.7* | *<8.7* | *5.7* | *6.7* | *8.7* | *9.9* |
| *Dicycloverine* | *<2.4* | *<2.1* | *<2.6* | *<2.3* | *2.1* | *2.6* | *2.3* | *2.6* |
| *Diltiazem* | *<2.5* | *<2.2* | *<2.5* | *<2.2* | *2.2* | *2.5* | *2.2* | *2.5* |
| *Diphenhydramine* | *<2.9* | *<2.7* | *<3* | *<8.3* | *2.7* | *3* | *8.3* | *9.4* |
| *Disopyramide* | *<3.7* | *<2.5* | *<4* | *<7* | *2.5* | *4* | *7* | *7.8* |
| *Donepezil* | *<1.7* | *<1.5* | *<1.8* | *<0.82* | *1.5* | *1.8* | *0.82* | *0.9* |
| *Erythromycin* | *<1.7* | *<1.8* | *<1.6* | *<4.5* | *1.6* | *1.8* | *4.5* | *5.1* |
| *Fenofibrate* | *<1.8* | *<1.8* | *<2.5* | *<8.6* | *1.8* | *2.5* | *8.6* | *9.7* |
| ***Fexofenadine*** | ***<2.3*** | ***<2.1*** | ***<2.4*** | ***<9.5*** | ***2.1*** | ***2.4*** | ***9.5*** | ***11*** |
| *Glibenclamide* | *<2* | *<1.9* | *<2.1* | *<8.7* | *1.9* | *2.1* | *8.7* | *9.8* |
| *Glimepiride* | *<2.3* | *<2.2* | *<2.4* | *<8.5* | *2.2* | *2.4* | *8.5* | *9.6* |
| *Haloperidol* | *<2.9* | *<2.5* | *<2.8* | *<1.5* | *2.5* | *2.9* | *1.5* | *1.7* |
| *Iopromide* | *<15* | *<12* | *<16* | *<12* | *12* | *16* | *12* | *12* |
|  | **Horni Machovec** | **Cep** | **Zlata Stoka** | **Černiš** | **LOQ_1_ min** | **LOQ_1_ max** | **LOQ_2_ min** | **LOQ_2_ max** |
| *Irbesartan* | *<2.2* | *<2* | *<2.3* | *<7.8* | *2* | *2.3* | *7.8* | *8.8* |
| *Lamotrigine* | *<11* | *<10* | *<11* | *<10* | *10* | *11* | *10* | *11* |
| *Loperamide* | *<2.3* | *<2.2* | *<2.4* | *<2.3* | *2.2* | *2.4* | *2.3* | *2.6* |
| *Maprotiline* | *<2.6* | *<2.5* | *<2.8* | *<8.5* | *2.5* | *2.8* | *8.5* | *9.6* |
| *Meclozine* | *<3.1* | *<3.9* | *<4* | *<1.5* | *3.1* | *4* | *1.5* | *1.7* |
| *Memantine* | *<3.4* | *<3* | *<3.7* | *<7.9* | *3* | *3.7* | *7.9* | *8.7* |
| ***Metoprolol*** | ***<2.2*** | ***<2.1*** | ***<2.5*** | ***<1.8*** | ***2.1*** | ***2.5*** | ***1.8*** | ***1.9*** |
| ***Metoprolol acid*** | ***<1.2*** | ***<0.95*** | ***13*** | ***<8.4*** | ***0.95*** | ***1.2*** | ***8.4*** | ***8.6*** |
| *Mianserin* | *<2.2* | *<2.1* | *<2.3* | *<2.3* | *2.1* | *2.3* | *2.3* | *2.6* |
| *Miconazole* | *<1.6* | *<2.2* | *<2.5* | *<1.2* | *1.6* | *2.5* | *1.2* | *1.3* |
| *Mirtazapine* | *<4* | *<2.5* | *<4.4* | *43* | *2.5* | *4.4* | *4.7* | *5.2* |
| *Orphenadrine* | *<1.2* | *<1.2* | *<1.3* | *<8.5* | *1.2* | *1.3* | *8.5* | *9.6* |
| *Oseltamivir carboxylate* | *<3.2* | *<2.4* | *<3.3* | *<8.1* | *2.4* | *3.3* | *8.1* | *8.5* |
| *Oxazepam* | *<1.4* | *<1.4* | *<1.9* | *<0.93* | *1.4* | *1.9* | *0.93* | *1.1* |
| *Oxcarbazepine* | *<3.4* | *<2.5* | *<4.5* | *<9* | *2.5* | *4.5* | *9* | *10* |
| *Pizotifen* | *<2.9* | *<2.1* | *<3* | *<8.7* | *2.1* | *3* | *8.7* | *9.8* |
| *Propranolol* | *<1.8* | *<1.7* | *<2* | *<1.5* | *1.7* | *2* | *1.5* | *1.7* |
| *Ropinirole* | *<3.3* | *<2.5* | *<2.9* | *<1.4* | *2.5* | *3.3* | *1.4* | *1.5* |
| *Rosuvastatin* | *<0.9* | *<0.82* | *<0.9* | *<7.5* | *0.82* | *0.9* | *7.5* | *8.4* |
| *Roxythromycin* | *<2.3* | *<2.2* | *<2.4* | *<4.9* | *2.2* | *2.4* | *4.9* | *5.3* |
| *Sertraline* | *<2.2* | *<2.1* | *<2.3* | *<2* | *2.1* | *2.3* | *2* | *2.3* |
| *Norsertraline* | *<5.6* | *<6* | *<7.3* | *<2* | *5.6* | *7.3* | *2* | *2.3* |
| *Sotalol* | *<36* | *<45* | *<47* | *<12* | *36* | *47* | *12* | *12* |
| *Sulfadiazine* | *<20* | *<12* | *<19* | *<11* | *12* | *20* | *11* | *12* |
| *Sulfamerazine* | *<5.2* | *<3.9* | *<4.7* | *<13* | *3.9* | *5.2* | *13* | *14* |
| *Sulfamethazine* | *<4.4* | *<3.3* | *<4* | *<14* | *3.3* | *4.4* | *14* | *15* |
| *Sulfamethizole* | *<4.9* | *<2.7* | *<4.4* | *<14* | *2.7* | *4.9* | *14* | *15* |
| ***Sulfamethoxazole*** | ***<5.8*** | ***<4.3*** | ***<5.2*** | ***<13*** | ***4.3*** | ***5.8*** | ***13*** | ***14*** |
| ***N1_Acetylsufamethaxazole*** | ***<16*** | ***<12*** | ***<15*** | ***<11*** | ***12*** | ***16*** | ***11*** | ***12*** |
| ***N4_Acetylsufamethaxazole*** | ***<9.2*** | ***<6.9*** | ***<11*** | ***<12*** | ***6.9*** | ***11*** | ***12*** | ***12*** |
| *Sulfapyridine* | *<0.82* | *<0.66* | *<0.78* | *<11* | *0.66* | *0.82* | *11* | *12* |
| *Tamoxifen* | *<1* | *<0.95* | *<1* | *<1.8* | *0.95* | *1* | *1.8* | *2* |
| ***Telmisartan*** | ***<3.2*** | ***<3.1*** | ***5.6*** | ***<19*** | ***3.1*** | ***3.8*** | ***19*** | ***23*** |
| *Terbinafine* | *<2.1* | *<2* | *<2.2* | *<7.4* | *2* | *2.2* | *7.4* | *8.3* |
| *Terbutaline* | *<8.2* | *<7.8* | *<8.6* | *<11* | *7.8* | *8.6* | *11* | *12* |
| *Theophylline* | *<8.9* | *<7.3* | *<8.2* | *10* | *7.3* | *8.9* | *7.6* | *7.9* |
| ***Tramadol*** | ***<11*** | ***<9.9*** | ***14*** | ***<9.3*** | ***9.9*** | ***12*** | ***9.3*** | ***10*** |
| *Trazodone* | *<2.3* | *<1.9* | *<2.2* | *<2* | *1.9* | *2.3* | *2* | *2.2* |
| *Triamterene* | *<4* | *<4.4* | *<4.2* | *<7* | *4* | *4.4* | *7* | *7.2* |
| ***Trimethoprim*** | ***<4.2*** | ***<4*** | ***<4.2*** | ***<8.7*** | ***4*** | ***4.2*** | ***8.7*** | ***9.2*** |
| ***Valsartan*** | ***<4.1*** | ***<4.2*** | ***15*** | ***<7*** | ***4.1*** | ***4.6*** | ***7*** | ***7.9*** |
| ***Venlafaxine*** | ***<2.6*** | ***<2.4*** | ***4*** | ***<6.8*** | ***2.4*** | ***2.9*** | ***6.8*** | ***7.6*** |
| ***O-Desmethylvenlafaxine*** | ***<2.2*** | ***<1.6*** | ***10*** | ***<7.5*** | ***1.6*** | ***2.2*** | ***7.5*** | ***8.4*** |
| *Verapamil* | *<1.7* | *<1.6* | *<1.8* | *3.5* | *1.6* | *1.8* | *2.1* | *2.4* |
| *Vortioxetine* | *<2.4* | *<2.3* | *<3.9* | *<1.6* | *2.3* | *3.9* | *1.6* | *1.8* |

**Table S3.3.** **Pesticide concentrations in the sampling sites and the minimum and maximum LOQ values (ng.L^-1^).** LOQ_1_ = LOQ values for Horni Machovec, Cep and Zlata stoka sampling sites; LOQ_2_ = LOQ values for Černiš site.

|  | | **Horni Machovec** | | **Cep** | **Zlata Stoka** | | **Černiš** | | **LOQ_1_ min** | | **LOQ_1_ max** | | **LOQ_2_ min** | | **LOQ_2_ max** | |
| --- | --- | --- | --- | --- | --- | --- | --- | --- | --- | --- | --- | --- | --- | --- | --- | --- |
| 1-(3.4-Dichlorophenyl)_urea | | <16 | | <24 | <27 | | <3.4 | | 16 | | 27 | | 3.4 | | 4 | |
| 1H-benzotriazol | | <51 | | <69 | <58 | | <43 | | 51 | | 69 | | 43 | | 51 | |
| 1H-benzotriazol_(5/4)-methyl | | <6.2 | | <7.6 | 19 | | <7.7 | | 6.2 | | 7.6 | | 7.7 | | 9 | |
| 1H-benzotriazol_1-methyl | | <3.1 | | <5.2 | <5 | | <8.4 | | 3.1 | | 5.2 | | 8.4 | | 9.9 | |
| 2.4.5-trichlorophenoxyacetic_acid | | <2.3 | | <2.5 | <3.2 | | <4.7 | | 2.3 | | 3.2 | | 4.7 | | 5.4 | |
| 2.4-D | | <2.5 | | <2.3 | <3.5 | | <5.7 | | 2.3 | | 3.5 | | 5.7 | | 6.6 | |
| 2.4-Dichlorphenoxypropionic_acid | | <1.7 | | <1.8 | <2.4 | | <5.4 | | 1.7 | | 2.4 | | 5.4 | | 6.2 | |
| 3-chloro-4-methylaniline | | <2 | | <2.3 | <3.1 | | <8.2 | | 2 | | 3.1 | | 8.2 | | 9.6 | |
| 4-Isopropylaniline | | <6.2 | | <11 | <23 | | <4 | | 6.2 | | 23 | | 4 | | 4.7 | |
| Acetochlor | | <4.4 | | <5 | <4.6 | | <3 | | 4.4 | | 5 | | 3 | | 3.6 | |
| Acetochlor_ESA | | 24 | | <3.7 | 14 | | 270 | | 3.7 | | 6 | | 5.4 | | 6.2 | |
| Acetochlor_OA | | <29 | | <22 | <38 | | 45 | | 22 | | 38 | | 4.5 | | 5.2 | |
| Alachlor | | <1.2 | | <1.4 | <0.99 | | <3 | | 0.99 | | 1.4 | | 3 | | 3.6 | |
| Alachlor_ESA | | <32 | | <23 | 41 | | 720 | | 23 | | 33 | | 20 | | 24 | |
| Alachlor_OA | | 9.4 | | <4.6 | <5.6 | | 22 | | 4.6 | | 5.6 | | 6.2 | | 7.2 | |
| Ametryn | | <3.6 | | <5 | <3.3 | | <3.2 | | 3.3 | | 5 | | 3.2 | | 3.8 | |
| Anthranilic_acid_isopropylamide | | <2.1 | | <3.2 | <2.7 | | <3.9 | | 2.1 | | 3.2 | | 3.9 | | 4.6 | |
| Atraton | | <3.3 | | <3.6 | <3 | | <3.8 | | 3 | | 3.6 | | 3.8 | | 4.5 | |
| Atrazine | | <2.5 | | <2.9 | <2.4 | | <3.5 | | 2.4 | | 2.9 | | 3.5 | | 4.1 | |
| Atrazine_2-hydroxy | | 27 | | <3.1 | 11 | | 62 | | 2.8 | | 3.1 | | 1.2 | | 1.4 | |
| Atrazine_desethyl | | <5.5 | | <8.2 | <6.9 | | <4.1 | | 5.5 | | 8.2 | | 4.1 | | 4.8 | |
| Atrazine_desethyl-2-hydroxy | | <3.4 | | <3.6 | <2.7 | | 15 | | 2.7 | | 3.6 | | 6.2 | | 7.3 | |
| Atrazine_desisopropyl | | <6.2 | | <10 | <8.9 | | <4 | | 6.2 | | 10 | | 4 | | 4.7 | |
| Azoxystrobin | | <9.6 | | <12 | <7.7 | | <0.84 | | 7.7 | | 12 | | 0.84 | | 1 | |
| Bensulfuron_methyl | | <4.8 | | <5.6 | <4.4 | | <0.78 | | 4.4 | | 5.6 | | 0.78 | | 0.92 | |
| Bentazone | | <0.39 | | <0.44 | 2.5 | | <6.2 | | 0.39 | | 0.54 | | 6.2 | | 7.1 | |
| Carbendazim | | <3.3 | | <4.6 | <2.9 | | <4.1 | | 2.9 | | 4.6 | | 4.1 | | 4.8 | |
| Carbofuran-3-hydroxy | | <4.9 | | <7.3 | <6.1 | | <4.3 | | 4.9 | | 7.3 | | 4.3 | | 5.1 | |
| Clomazone | | <4.5 | | <4.9 | <4.4 | | <3.1 | | 4.4 | | 4.9 | | 3.1 | | 3.6 | |
| Cyanazine | | <8.3 | | <9.8 | <8.2 | | <3.8 | | 8.2 | | 9.8 | | 3.8 | | 4.5 | |
| Cyproconazole | | 2.9 | | <1.7 | <1.6 | | <0.81 | | 1.5 | | 1.7 | | 0.81 | | 0.96 | |
| DEET | | 49000 | | 1300 | 1200 | | 1100 | | 23 | | 34 | | 39 | | 46 | |
| Desmetryn | | <4.3 | | <5.2 | <3.4 | | <3.3 | | 3.4 | | 5.2 | | 3.3 | | 3.9 | |
| Diazinon | | <6.6 | | <7.6 | <7 | | <0.87 | | 6.6 | | 7.6 | | 0.87 | | 1 | |
| Dimethachlor | | <6.2 | | <7.1 | <6.5 | | <3.4 | | 6.2 | | 7.1 | | 3.4 | | 4 | |
| Dimethachlor_ESA | | <6.4 | | <4.2 | 8.4 | | 13 | | 4.2 | | 6.4 | | 3.8 | | 4.5 | |
| Dimethachlor_OA | | <4.8 | | <4.8 | <4 | | <3.9 | | 4 | | 4.8 | | 3.9 | | 4.6 | |
| Dimethenamid_ESA | | <10 | | <8.3 | <8.3 | | <3.8 | | 8.3 | | 10 | | 3.8 | | 4.5 | |
| Dimethenamid_OA | | <12 | | <13 | <11 | | <6.3 | | 11 | | 13 | | 6.3 | | 7.4 | |
| Dimethoate | | <11 | | <11 | <11 | | <4 | | 11 | | 11 | | 4 | | 4.7 | |
|  | **Horni Machovec** | | **Cep** | | | **Zlata Stoka** | | **Černiš** | | **LOQ_1_ min** | | **LOQ_1_ max** | | **LOQ_2_ min** | | **LOQ_2_ max** |
| Diuron | 2.3 | | 2.5 | | | <1.7 | | <3.8 | | 1.4 | | 2.1 | | 3.8 | | 4.4 |
| Diuron_desmethyl | <6.1 | | 11 | | | <7.5 | | <3.6 | | 6.1 | | 9.1 | | 3.6 | | 4.2 |
| Epoxiconazole | <5.8 | | <6.6 | | | <6.1 | | <0.81 | | 5.8 | | 6.6 | | 0.81 | | 0.96 |
| Fenuron | <6.4 | | <9.3 | | | <8 | | <3.7 | | 6.4 | | 9.3 | | 3.7 | | 4.3 |
| Florasulam | <6.2 | | <9.2 | | | <7.7 | | <3.7 | | 6.2 | | 9.2 | | 3.7 | | 4.4 |
| Fluazifop-p | <5 | | <4.7 | | | <4.3 | | <0.86 | | 4.3 | | 5 | | 0.86 | | 1 |
| Flusilazole | <5.9 | | <6.8 | | | <6.2 | | <2.4 | | 5.9 | | 6.8 | | 2.4 | | 2.9 |
| Foramsulfuron | <8.7 | | <13 | | | <11 | | <41 | | 8.7 | | 13 | | 41 | | 49 |
| Hexazinone | <12 | | <15 | | | <12 | | <1 | | 12 | | 15 | | 1 | | 1.2 |
| Chlorantraniliprole | <1.8 | | <2.1 | | | <1.9 | | <0.8 | | 1.8 | | 2.1 | | 0.8 | | 0.95 |
| Chloridazon | <14 | | <16 | | | <14 | | <4 | | 14 | | 16 | | 4 | | 4.7 |
| Chloridazon_desphenyl | <210 | | <140 | | | <88 | | <37 | | 88 | | 210 | | 37 | | 44 |
| Chloridazon_methyl_desphenyl | <19 | | <11 | | | <7.2 | | <3.6 | | 7.2 | | 19 | | 3.6 | | 4.2 |
| Chlorotoluron | <9.6 | | <11 | | | <9.4 | | <3.7 | | 9.4 | | 11 | | 3.7 | | 4.4 |
| Chlorotoluron_desmethyl | <4.7 | | <7 | | | <5.8 | | <3.8 | | 4.7 | | 7 | | 3.8 | | 4.5 |
| Chlorpyrifos | <68 | | <78 | | | <72 | | <5.9 | | 68 | | 78 | | 5.9 | | 7 |
| Imazamethabenz_methyl | <5.7 | | <8.4 | | | <7 | | <3.4 | | 5.7 | | 8.4 | | 3.4 | | 3.9 |
| Imazamox | <6.3 | | <7.4 | | | <5.2 | | <3.9 | | 5.2 | | 7.4 | | 3.9 | | 4.6 |
| Imidacloprid | <12 | | <13 | | | <11 | | <3.2 | | 11 | | 13 | | 3.2 | | 3.8 |
| Ioxynil | <2.3 | | <2.6 | | | <3.4 | | <1.5 | | 2.3 | | 3.4 | | 1.5 | | 1.7 |
| Isoproturon | <7.4 | | <11 | | | <9.2 | | <3.8 | | 7.4 | | 11 | | 3.8 | | 4.4 |
| Isoproturon_monodemethyl | <6 | | <8.9 | | | <7.4 | | <3.7 | | 6 | | 8.9 | | 3.7 | | 4.3 |
| Lenacil | <6.1 | | <7 | | | <5.9 | | <3.3 | | 5.9 | | 7 | | 3.3 | | 3.9 |
| Linuron | <3.6 | | <4 | | | <3.8 | | <0.85 | | 3.6 | | 4 | | 0.85 | | 1 |
| Malathion | <8.7 | | <10 | | | <9.2 | | <3 | | 8.7 | | 10 | | 3 | | 3.5 |
| MCPA | <0.73 | | <0.75 | | | <1.1 | | <0.73 | | 0.57 | | 1.1 | | 0.73 | | 0.83 |
| MCPP | <1.6 | | <1.7 | | | <2.3 | | <5.3 | | 1.2 | | 2.3 | | 5.3 | | 6 |
| Metalaxyl | <12 | | <17 | | | <14 | | <3.5 | | 12 | | 17 | | 3.5 | | 4.1 |
| Metazachlor | <6.3 | | 8.6 | | | 7.8 | | <3.3 | | 6.3 | | 7.3 | | 3.3 | | 3.9 |
| Metazachlor_ESA | <30 | | <26 | | | <38 | | <6.7 | | 26 | | 38 | | 6.7 | | 7.7 |
| Metazachlor_OA | 12 | | <12 | | | 25 | | 6.3 | | 11 | | 12 | | 4 | | 4.7 |
| Metconazole | <5.3 | | <6.1 | | | <4.3 | | <0.86 | | 4.3 | | 6.1 | | 0.86 | | 1 |
| Methabenzthiazuron | <7.3 | | <11 | | | <9 | | <3.7 | | 7.3 | | 11 | | 3.7 | | 4.3 |
| Methoxyfenozide | <6.4 | | <7.4 | | | <6.8 | | <0.88 | | 6.4 | | 7.4 | | 0.88 | | 1 |
| Metobromuron | <7.6 | | <11 | | | <9.4 | | <4.4 | | 7.6 | | 11 | | 4.4 | | 5.2 |
| Metolachlor | <6.1 | | <7 | | | <6.5 | | <3.7 | | 6.1 | | 7 | | 3.7 | | 4.3 |
| Metolachlor_ESA | 99 | | <13 | | | 23 | | 97 | | 12 | | 15 | | 3.5 | | 4.1 |
| Metolachlor_OA | <11 | | <15 | | | <12 | | 5.9 | | 11 | | 15 | | 3.5 | | 4.1 |
| Metoxuron | <14 | | <18 | | | <14 | | <3.8 | | 14 | | 18 | | 3.8 | | 4.5 |
| Metribuzin | <34 | | <45 | | | <37 | | <42 | | 34 | | 45 | | 42 | | 50 |
| Metribuzin_desamino | <5.3 | | <7.8 | | | <6.5 | | <3.5 | | 5.3 | | 7.8 | | 3.5 | | 4.1 |
| Metsulfuron_methyl | <4.6 | | <6.5 | | | <4.9 | | <3.6 | | 4.6 | | 6.5 | | 3.6 | | 4.2 |
| Monolinuron | 3.3 | | 3.4 | | | <2.8 | | <0.74 | | 2.1 | | 2.9 | | 0.74 | | 0.87 |
| N-chloroacetyl-2.6-diethylaniline | <8 | | <12 | | | <9.8 | | <0.43 | | 8 | | 12 | | 0.43 | | 0.51 |
| Picloram | <84 | | <71 | | | <36 | | <440 | | 36 | | 84 | | 440 | | 520 |
| Pirimicarb | <6 | | <6.2 | | | <5.7 | | <1.2 | | 5.7 | | 6.2 | | 1.2 | | 1.3 |
|  | **Horni Machovec** | | **Cep** | | | **Zlata Stoka** | | **Černiš** | | **LOQ_1_ min** | | **LOQ_1_ max** | | **LOQ_2_ min** | | **LOQ_2_ max** |
| Pirimiphos_methyl | <5.9 | | <6.8 | | | <6.2 | | 1.4 | | 5.9 | | 6.8 | | 0.86 | | 1 |
| Prometryn | <9.2 | | <14 | | | <8.9 | | <1 | | 8.9 | | 14 | | 1 | | 1.2 |
| Propachlor | <4.8 | | <5.6 | | | <6.2 | | <3.2 | | 4.8 | | 6.2 | | 3.2 | | 3.8 |
| Propazine | <5.7 | | <6.6 | | | <6 | | <0.86 | | 5.7 | | 6.6 | | 0.86 | | 1 |
| Propazine_2-hydroxy | <5.3 | | <7.6 | | | <6.3 | | 4.4 | | 5.1 | | 7.6 | | 3.8 | | 4.4 |
| Propiconazole | <5.4 | | <6.2 | | | <5.7 | | <0.83 | | 5.4 | | 6.2 | | 0.83 | | 0.99 |
| Pyrimethanil | <5.9 | | <8.8 | | | <7.3 | | <3.5 | | 5.9 | | 8.8 | | 3.5 | | 4.2 |
| Sebuthylazine | <5.8 | | <6.7 | | | <6.2 | | <3 | | 5.8 | | 6.7 | | 3 | | 3.6 |
| Simazine | <4.7 | | <7 | | | <5.8 | | <0.98 | | 4.7 | | 7 | | 0.98 | | 1.2 |
| Simazine_hydroxy | <8.3 | | <12 | | | <7.3 | | <3.8 | | 7.3 | | 12 | | 3.8 | | 4.4 |
| Tebuconazole | <5.3 | | <6.2 | | | 10 | | <0.55 | | 5.3 | | 6.2 | | 0.55 | | 0.65 |
| Terbuthylazine | <8.7 | | <10 | | | <9.1 | | 6.2 | | 8.7 | | 10 | | 3.8 | | 4.5 |
| Terbuthylazine_desethyl | 9.3 | | 13 | | | <8.2 | | <3.5 | | 6.6 | | 9.8 | | 3.5 | | 4.1 |
| Terbuthylazine_desethyl-2-hydroxy | <7 | | <11 | | | <5.8 | | 5.2 | | 5.8 | | 11 | | 1 | | 1.2 |
| Terbuthylazine_hydroxy | 150 | | <7.1 | | | 36 | | 13 | | 4.8 | | 7.1 | | 3.8 | | 4.5 |
| Terbutryn | <7.2 | | <11 | | | <8.9 | | <1 | | 7.2 | | 11 | | 1 | | 1.2 |
| Thiamethoxam | <6.8 | | <6.4 | | | <6.1 | | <3.8 | | 6.1 | | 6.8 | | 3.8 | | 4.5 |
| Triadimenol | <5.2 | | <5 | | | <4.3 | | <2.3 | | 4.3 | | 5.2 | | 2.3 | | 2.7 |
| Triallat | <11 | | <12 | | | <11 | | <3.1 | | 11 | | 12 | | 3.1 | | 3.7 |
| Triticonazole | <4.9 | | <5.7 | | | <4.6 | | <0.69 | | 4.6 | | 5.7 | | 0.69 | | 0.82 |
| Warfarin | <9.3 | | <8.5 | | | <7.6 | | <0.76 | | 7.6 | | 9.3 | | 0.76 | | 0.9 |

**Supplementary Information S4: Data collection and analysis**

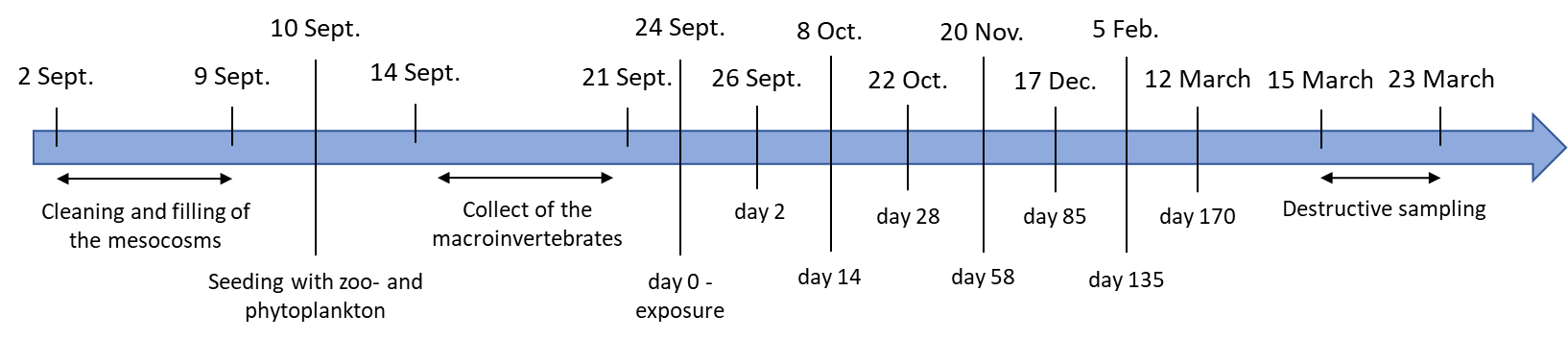

**Figure S4.1.** Timeline of the winter experiment in 2020-2021

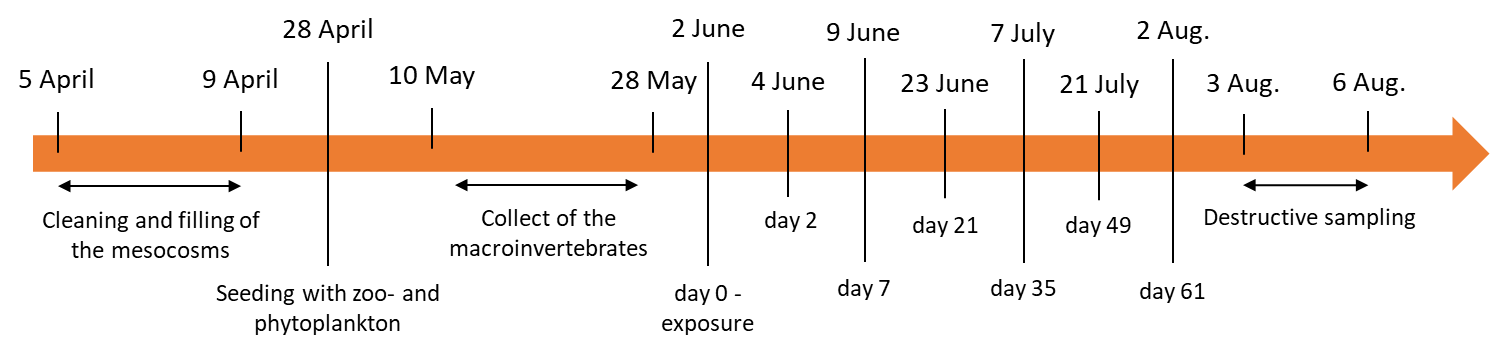

**Figure S4.2**. Timeline of the summer experiment in 2021

**Table S4.1.** **Response variables collected during the experiments.** This paper analyses only variables collected in the water column (‘pelagic’).

| **Response variable** | **Pelagic /**  **repeated** | **Pelagic/ start and end** | **Benthic** |
| --- | --- | --- | --- |
| Environmental conditions | X | - | - |
| Chlorophyll-*a* concentration | X | - | - |
| Zooplankton abundance | X | - | - |
| Chemical analysis | - | X | - |
| Predator emergence (Odonata) | X | - | - |
| Predator abundance (Odonata and *Notonecta*) | - | X | - |
| Other macroinvertebrate abundance | - | - | X |
| Other insect emergence | - | - | X |
| Leaf litter (decomposition bags) | - | - | X |
| Phytoplankton community composition and abundance | - | X (end) | - |
| Periphyton community composition and abundance | - | - | X |

**Table S4.2. Estimated individual dry weights for the zooplankton taxa and wet weights for the insect predators found in the mesocosms.**

| Taxon | Weight (µg) | Source |
| --- | --- | --- |
| Copepoda | 15 | Dumont et al. 1975 |
| *Chydorus* sp. | 2.5 | Dumont et al. 1975 |
| *Bosmina* sp | 3.8 | Dumont et al. 1975 |
| *Scapholeberis* sp. | 6.0 | Dumont et al. 1975 |
| *Ceriodaphnia* sp. | 6.3 | Dumont et al. 1975 |
| *Daphnia* sp. | 102 | Dumont et al. 1975 |
| Ostracoda | 15 | Widbom 1984 |
| Odonata | 2 × 10^5^ | Own measurements |
| *Notonecta* sp. | 2 × 10^5^ | Own measurements |

**S4.1. Processing of the phytoplankton samples**

After complete sedimentation in the laboratory, the water was decanted, and the sample was transferred to a Bürker counting chamber. Algal cells were counted using an optical microscope (OLYMPUS CX23) at 400× magnification. Taxa were identified to species or genus level (where possible) according to Kaštovský et al. (2018a, 2018b). We used the currently accepted phytoplankton nomenclature based on the AlgaeBase (Guiry and Guiry, 2023).

**S4.2. Details of statistical analyses**

Temporal changes in environmental parameters (pH, dissolved O_2_, conductivity, turbidity, and water level) and chlorophyll-a concentrations during the experiment were tested by a redundancy analysis (RDA on centred data) separately for the summer and the winter experiment. Treatment was used as a categorical explanatory variable and environmental variables were used as a response. “Day of the experiment” was used as a covariate. The statistical significance of the RDAs was tested by Monte Carlo permutation tests (999 unrestricted permutations under the reduced model in split-plot hierarchical design).

Differences in community composition of algal groups and taxa between treatments were visualised by principal component analyses (PCAs) with centred and log-transformed abundance data. The analysis was run on data collected at the end of each experiment and only species found in more than three samples in the given experiment were included in the analysis.

Changes in the zooplankton community structure were analysed using the principal response curve (PRC) method (Van den Brink and ter Braak, 1999) using the most practical level of taxonomic identification for each group. Interaction of treatment and day was used as the explanatory variable and community composition as a response. “Day of the experiment” was used as a covariate. Before analysis, the biomass of each taxon was log-transformed. Monte Carlo permutation tests were performed on each sampling date to identify the dates on which the treatments differed significantly. In all analyses, the permutations were set in a hierarchical split-plot design and thus permuted for each mesocosm individually. All multivariate and PRC analyses were conducted using CANOCO 5 (ter Braak and Smilauer, 2012).

Log-link Gamma GLMs were used for the analyses of the treatment effects on nutrients (dissolved organic carbon, nitrogen and phosphorus concentrations). Binomial GLMs were used for the analyses of treatment effects on cumulative emergence of predators in summer and on predator survival in winter, respectively. These analyses used only data from the end of each experiment. The probability of successful emergence of odonates in the summer experiment was defined as the ratio of the estimated number of individuals emerged by Day 56 to the size of the inoculum. Similarly, survival probability of odonates and *Notonecta* in the winter experiment was defined as the ratio of the estimated number of individuals collected on Day 169 to the size of the inoculum.

Gamma (copepods in winter) and zero-inflated Gamma GLMMs (*Daphnia* in summer and winter, and copepods in summer) with log link were used in the analyses of the treatment effects on *Daphnia* and copepod biomass dynamics. Zero inflation was included where necessary to account for the presence of zeroes in the data. Preliminary inspection of the time series suggested unidirectional or unimodal changes in all response variables. These trends were approximated by first- and second-order polynomial dependence on time, and different shapes of this dependence were described phenomenologically by expressing time in days, days^0.5^ and days^0.25^ from the start of the experiment as in Yildiz et al. (2021). The candidate model set thus contained 12 models for responses measured during the summer and winter experiments (3 shapes of temporal dependence x linear or quadratic dependence on time x presence or absence of statistical interaction between temporal dependence and treatment; Table S7.1). The effect of treatment was then tested with the most parsimonious temporal structure identified in the preceding step, and the candidate model set contained 8 models for responses measured during the summer and winter experiments (Heating and pharmaceuticals alone or in combination, Table S7.2).

GLMMs were implemented using the ‘glmmTMB’ function from the *glmmTMB* package (Brooks et al., 2017). Model residuals were inspected in the *DHARMa* package (Hartig, 2020) to confirm the lack of overdispersion, zero inflation (where appropriate) and any remaining major trends. Parameter values of the most parsimonious (hereafter ‘best’) models are reported along with their 95% confidence interval, calculated with the *sjPlot* package (Lüdecke, 2020), and model results were visualized using the *ggeffects* package (Lüdecke, 2018). Model selection approach with the corrected Akaike information criterion was used to identify the most parsimonious model for each univariate response.

Finally, to disentangle the direct and indirect effects of the pharmaceutical mixture and warming on interactions within the aquatic community, we tested simple networks of cause-effect relationships describing a trophic cascade and predator-prey relationships using confirmatory path analysis (piecewise structural equation models [SEM]; Shipley and Douma, 2020), implemented in *piecewiseSEM* package (Lefcheck, 2016). This approach can identify mechanistic pathways underlying the effects of different stressors on aquatic ecosystems (Schmidt et al., 2022). Using *a priori* knowledge, we built a complete model of cause-effect relationships based on hypothetical pathways supported by literature (Table S1.1). In brief, we assumed that the direct effects of pharmaceuticals and warming on predators (odonates in summer, and odonates and *Notonecta* in winter) will indirectly affect filter feeder biomass (= cladocerans in our mesocosms) by top-down control. Pharmaceuticals and warming were also assumed to directly reduce the biomass of filter feeders and thus indirectly increase phytoplankton biomass due to reduced grazing pressure (Belden et al., 2007). Phytoplankton (chlorophyll-a concentration), filter feeder biomass and predator biomass were used as response variables. We used predator biomass as a proxy of their feeding pressure on lower trophic levels. In the summer experiment, we estimated it by the temporally averaged cumulative emergence of odonates as almost no larvae survived until the end of the experiment (note that almost all larvae emerged between Day 42 and Day 56 of the experiment or died). In the winter experiment, we used the survival of *Notonecta* and odonate larvae as no adults emerged during that experiment (C. Duchet et al., unpublished data). We used Fisher’s C statistics to assess the overall fit of each SEM model, with *p* > 0.05 as indicating a good fit. All these analyses were done in R version 4.1.2 (R Core Team, 2018).

**S5. Environmental parameters values**

**Table S5.1.** **Environmental variables measured at each sampling date** (mean ± SE, *n* = 7 for each treatment) **in the summer experiment.** CLT: non-heated, without pharmaceuticals; CHT: heated, without pharmaceuticals; TLT: non-heated, with pharmaceuticals; THT: heated, with pharmaceuticals.

| Variable |  | **Day of experiment** | | | | | | | | | | | | | | | | | | | | | | |
| --- | --- | --- | --- | --- | --- | --- | --- | --- | --- | --- | --- | --- | --- | --- | --- | --- | --- | --- | --- | --- | --- | --- | --- | --- |
|  | Treatment | 0 | | | 2 | | | 7 | | | | 22 | | | 35 | | | | 49 | | | 61 | | |
| Temperature (°C) | CLT | **14.9** | ± | 0.05 | **17.9** | ± | 0.36 | | **19.4** | ± | 0.13 | **21.9** | ± | 0.17 | **21.8** | ± | 0.13 | **20.3** | | ± | 0.07 | **19.9** | ± | 0.26 |
|  | CHT | **14.6** | ± | 0.20 | **21.7** | ± | 0.41 | | **23.0** | ± | 0.15 | **25.4** | ± | 0.38 | **25.4** | ± | 0.26 | **24.2** | | ± | 0.26 | **23.6** | ± | 0.24 |
|  | TLT | **14.7** | ± | 0.25 | **17.9** | ± | 0.31 | | **19.5** | ± | 0.30 | **21.9** | ± | 0.22 | **21.8** | ± | 0.18 | **20.2** | | ± | 0.10 | **20.1** | ± | 0.33 |
|  | THT | **14.5** | ± | 0.15 | **21.7** | ± | 0.49 | | **23.2** | ± | 0.24 | **25.5** | ± | 0.28 | **25.4** | ± | 0.10 | **24.3** | | ± | 0.28 | **23.8** | ± | 0.25 |
| pH | CLT | **8.43** | ± | 0.41 | **8.63** | ± | 0.41 | | **8.60** | ± | 0.36 | **8.04** | ± | 0.25 | **7.62** | ± | 0.05 | **7.56** | | ± | 0.05 | **7.54** | ± | 0.11 |
|  | CHT | **8.39** | ± | 0.26 | **8.43** | ± | 0.23 | | **8.36** | ± | 0.22 | **7.81** | ± | 0.21 | **7.56** | ± | 0.07 | **7.59** | | ± | 0.09 | **7.36** | ± | 0.28 |
|  | TLT | **8.41** | ± | 0.24 | **8.50** | ± | 0.17 | | **8.47** | ± | 0.13 | **7.92** | ± | 0.24 | **7.76** | ± | 0.12 | **7.65** | | ± | 0.05 | **7.68** | ± | 0.05 |
|  | THT | **8.40** | ± | 0.29 | **8.43** | ± | 0.22 | | **8.37** | ± | 0.28 | **7.72** | ± | 0.09 | **7.70** | ± | 0.08 | **7.67** | | ± | 0.13 | **7.60** | ± | 0.06 |
| Conductivity (mS.cm^-1^) | CLT | **229.2** | ± | 5.33 | **228.6** | ± | 5.04 | | **225.6** | ± | 6.00 | **222.9** | ± | 9.8 | **227.9** | ± | 5.81 | **229.4** | | ± | 4.35 | **230.4** | ± | 4.89 |
|  | CHT | **231.6** | ± | 4.39 | **233.6** | ± | 4.36 | | **235.7** | ± | 4.96 | **248.6** | ± | 4.5 | **251.6** | ± | 1.51 | **252.9** | | ± | 2.12 | **260.9** | ± | 2.85 |
|  | TLT | **227.2** | ± | 5.90 | **226.8** | ± | 6.42 | | **225.7** | ± | 5.71 | **226.8** | ± | 3.3 | **227.7** | ± | 2.69 | **227.6** | | ± | 4.54 | **227.4** | ± | 5.09 |
|  | THT | **231.0** | ± | 3.90 | **233.1** | ± | 3.37 | | **234.0** | ± | 3.96 | **245.6** | ± | 4.3 | **249.9** | ± | 1.86 | **251.6** | | ± | 2.15 | **257.6** | ± | 3.51 |
| Dissolved oxygen (mg.L^-1^) | CLT | **18.43** | ± | 1.83 | **9.65** | ± | 2.52 | | **15.09** | ± | 0.64 | **12.62** | ± | 0.43 | **12.78** | ± | 0.37 | **13.53** | | ± | 0.73 | **12.93** | ± | 1.15 |
|  | CHT | **18.64** | ± | 1.07 | **10.52** | ± | 3.62 | | **14.61** | ± | 0.68 | **12.51** | ± | 0.32 | **12.29** | ± | 0.27 | **12.91** | | ± | 0.57 | **13.22** | ± | 0.64 |
|  | TLT | **17.98** | ± | 0.81 | **16.21** | ± | 1.24 | | **15.20** | ± | 0.33 | **13.20** | ± | 0.41 | **13.05** | ± | 0.79 | **14.07** | | ± | 0.46 | **13.57** | ± | 0.44 |
|  | THT | **17.94** | ± | 0.53 | **16.48** | ± | 0.55 | | **14.77** | ± | 0.58 | **12.47** | ± | 0.41 | **12.61** | ± | 0.53 | **13.64** | | ± | 0.40 | **13.21** | ± | 0.46 |
| Turbidity (NTU) | CLT | **1.49** | ± | 0.23 | **1.69** | ± | 0.77 | | **1.61** | ± | 0.40 | **1.57** | ± | 0.41 | **1.23** | ± | 0.36 | **1.15** | | ± | 0.42 | **1.18** | ± | 0.45 |
|  | CHT | **1.62** | ± | 0.19 | **1.46** | ± | 0.26 | | **1.36** | ± | 0.18 | **1.30** | ± | 0.18 | **0.86** | ± | 0.22 | **0.94** | | ± | 0.26 | **0.94** | ± | 0.44 |
|  | TLT | **1.61** | ± | 0.17 | **1.45** | ± | 0.16 | | **1.47** | ± | 0.21 | **1.63** | ± | 0.39 | **1.05** | ± | 0.26 | **1.09** | | ± | 0.40 | **0.81** | ± | 0.21 |
|  | THT | **1.51** | ± | 0.21 | **1.22** | ± | 0.20 | | **1.26** | ± | 0.26 | **1.53** | ± | 0.34 | **0.92** | ± | 0.26 | **0.99** | | ± | 0.48 | **0.73** | ± | 0.22 |
| Water depth (cm) | CLT | **99.9** | ± | 0.65 | **99.4** | ± | 0.86 | | **99.3** | ± | 0.81 | **97.9** | ± | 0.92 | **99.1** | ± | 1.05 | **100.2** | | ± | 0.84 | **99.7** | ± | 1.16 |
|  | CHT | **99.9** | ± | 0.30 | **98.5** | ± | 0.59 | | **97.8** | ± | 0.45 | **93.5** | ± | 0.39 | **92.4** | ± | 0.43 | **90.4** | | ± | 0.43 | **87.2** | ± | 0.39 |
|  | TLT | **99.5** | ± | 0.35 | **98.9** | ± | 0.67 | | **98.7** | ± | 0.65 | **97.3** | ± | 1.60 | **99.2** | ± | 0.50 | **100.3** | | ± | 0.87 | **99.1** | ± | 1.12 |
|  | THT | **99.8** | ± | 0.65 | **98.8** | ± | 0.63 | | **98.2** | ± | 0.88 | **93.8** | ± | 0.65 | **92.4** | ± | 0.71 | **90.4** | | ± | 0.80 | **86.7** | ± | 0.62 |
| Chlorophyll-a  (RFU) | CLT | **0.22** | ± | 0.14 | **0.15** | ± | 0.06 | | **0.19** | ± | 0.06 | **1.06** | ± | 0.84 | **0.91** | ± | 0.54 | **0.85** | | ± | 0.56 | **0.80** | ± | 0.63 |
|  | CHT | **0.34** | ± | 0.20 | **0.20** | ± | 0.11 | | **0.38** | ± | 0.27 | **0.86** | ± | 0.56 | **1.01** | ± | 1.23 | **1.4** | | ± | 0.72 | **0.80** | ± | 0.74 |
|  | TLT | **0.21** | ± | 0.11 | **0.13** | ± | 0.07 | | **0.27** | ± | 0.09 | **0.86** | ± | 0.66 | **0.57** | ± | 0.25 | **0.53** | | ± | 0.14 | **0.75** | ± | 0.64 |
|  | THT | **0.34** | ± | 0.15 | **0.12** | ± | 0.06 | | **0.23** | ± | 0.06 | **1.09** | ± | 0.37 | **0.56** | ± | 0.23 | **1.24** | | ± | 0.96 | **0.66** | ± | 0.40 |

**Table S5.2: Environmental variables measured at each sampling date** (mean ± SE, *n* = 7 for each treatment) **in the winter experiment.** CLT: non-heated, without pharmaceuticals; CHT: heated, without pharmaceuticals; TLT: non-heated, with pharmaceuticals; THT: heated, with pharmaceuticals.

| Variable |  |  | | | | **Day of experiment** | | | | | | | | | | | | | | | | | | | | | | | | | | | | |
| --- | --- | --- | --- | --- | --- | --- | --- | --- | --- | --- | --- | --- | --- | --- | --- | --- | --- | --- | --- | --- | --- | --- | --- | --- | --- | --- | --- | --- | --- | --- | --- | --- | --- | --- |
|  | Treatment | | 0 | | | | 2 | | | | 14 | | | | 28 | | | | | 57 | | | | 84 | | | | 134 | | | | 169 | | |
| Temperature (°C) | CLT | **16.8** | | ± | 0.1 | | | **13.8** | ± | 0.1 | | **12.4** | ± | 0.1 | | **9.4** | ± | 0.2 | **7.0** | | ± | 0.2 | **3.5** | | ± | 0.3 | **5.0** | | ± | 1.4 | **3.9** | | ± | 0.1 |
|  | CHT | **20.8** | | ± | 0.2 | | | **17.6** | ± | 0.1 | | **16.2** | ± | 0.1 | | **13.0** | ± | 0.2 | **10.8** | | ± | 0.2 | **7.4** | | ± | 0.3 | **7.7** | | ± | 1.7 | **8.4** | | ± | 0.7 |
|  | TLT | **16.8** | | ± | 0.2 | | | **13.7** | ± | 0.2 | | **12.3** | ± | 0.2 | | **9.4** | ± | 0.1 | **6.9** | | ± | 0.1 | **3.5** | | ± | 0.2 | **4.4** | | ± | 0.2 | **4.0** | | ± | 0.1 |
|  | THT | **20.9** | | ± | 0.6 | | | **18.2** | ± | 1.2 | | **16.3** | ± | 0.2 | | **13.1** | ± | 0.2 | **10.8** | | ± | 0.1 | **7.1** | | ± | 0.1 | **8.5** | | ± | 0.5 | **8.2** | | ± | 0.3 |
| pH | CLT | **7.83** | | ± | 0.09 | | | **7.86** | ± | 0.10 | | **7.9** | ± | 0.27 | | **8.48** | ± | 0.21 | **8.71** | | ± | 0.33 | **8.34** | | ± | 0.17 | **8.58** | | ± | 0.23 | **8.36** | | ± | 0.15 |
|  | CHT | **7.79** | | ± | 0.16 | | | **7.76** | ± | 0.10 | | **7.8** | ± | 0.17 | | **8.29** | ± | 0.28 | **8.35** | | ± | 0.36 | **8.16** | | ± | 0.20 | **8.48** | | ± | 0.16 | **8.31** | | ± | 0.27 |
|  | TLT | **7.78** | | ± | 0.07 | | | **7.82** | ± | 0.09 | | **7.9** | ± | 0.21 | | **8.38** | ± | 0.17 | **8.49** | | ± | 0.14 | **8.19** | | ± | 0.02 | **8.68** | | ± | 0.17 | **8.51** | | ± | 0.33 |
|  | THT | **7.81** | | ± | 0.10 | | | **7.76** | ± | 0.09 | | **7.8** | ± | 0.15 | | **8.44** | ± | 0.22 | **8.36** | | ± | 0.08 | **8.16** | | ± | 0.03 | **8.59** | | ± | 0.14 | **8.44** | | ± | 0.27 |
| Conductivity (mS.cm^-1^) | CLT | **244** | | ± | 5 | | | **236** | ± | 4 | | **234** | ± | 4 | | **226** | ± | 5 | **217** | | ± | 8 | **219** | | ± | 8 | **217** | | ± | 16 | **208** | | ± | 9.25 |
|  | CHT | **244** | | ± | 4 | | | **238** | ± | 4 | | **239** | ± | 3 | | **237** | ± | 3 | **236** | | ± | 4 | **242** | | ± | 3 | **238** | | ± | 12 | **247** | | ± | 4.81 |
|  | TLT | **246** | | ± | 5 | | | **239** | ± | 5 | | **237** | ± | 5 | | **230** | ± | 5 | **223** | | ± | 6 | **224** | | ± | 6 | **217** | | ± | 7 | **213** | | ± | 6.26 |
|  | THT | **244** | | ± | 2 | | | **238** | ± | 2 | | **241** | ± | 2 | | **237** | ± | 2 | **236** | | ± | 2 | **243** | | ± | 2 | **243** | | ± | 4 | **247** | | ± | 4.89 |
| Dissolved oxygen (mg.L^-1^) | CLT | **8.79** | | ± | 0.28 | | | **9.55** | ± | 0.32 | | **11.87** | ± | 1.05 | | **14.86** | ± | 0.77 | **15.33** | | ± | 0.5 | **16.00** | | ± | 0.56 | **16.83** | | ± | 1.31 | **17.29** | | ± | 0.34 |
|  | CHT | **8.55** | | ± | 0.37 | | | **8.84** | ± | 0.38 | | **12** | ± | 0.33 | | **13.83** | ± | 0.75 | **14.22** | | ± | 0.68 | **15.05** | | ± | 0.5 | **15.32** | | ± | 1.21 | **15.92** | | ± | 0.66 |
|  | TLT | **8.41** | | ± | 0.21 | | | **9.42** | ± | 0.39 | | **12.54** | ± | 0.51 | | **14.38** | ± | 0.48 | **15.18** | | ± | 0.69 | **15.69** | | ± | 0.46 | **18.06** | | ± | 0.6 | **17.16** | | ± | 0.44 |
|  | THT | **8.8** | | ± | 0.19 | | | **8.83** | ± | 0.38 | | **12.06** | ± | 1.04 | | **14.4** | ± | 0.45 | **14.66** | | ± | 0.36 | **15.79** | | ± | 0.38 | **15.46** | | ± | 0.52 | **16.27** | | ± | 0.63 |
| Turbidity (NTU) | CLT | **0.69** | | ± | 0.38 | | | **0.80** | ± | 0.49 | | **0.64** | ± | 0.35 | | **0.53** | ± | 0.13 | **0.49** | | ± | 0.11 | **0.52** | | ± | 0.09 | **0.55** | | ± | 0.11 | **0.68** | | ± | 0.17 |
|  | CHT | **0.47** | | ± | 0.15 | | | **0.43** | ± | 0.09 | | **0.48** | ± | 0.11 | | **0.53** | ± | 0.14 | **0.49** | | ± | 0.05 | **0.55** | | ± | 0.13 | **0.48** | | ± | 0.11 | **0.58** | | ± | 0.09 |
|  | TLT | **0.67** | | ± | 0.19 | | | **0.70** | ± | 0.52 | | **0.62** | ± | 0.34 | | **0.54** | ± | 0.11 | **0.51** | | ± | 0.10 | **0.51** | | ± | 0.08 | **0.53** | | ± | 0.08 | **0.55** | | ± | 0.11 |
|  | THT | **0.67** | | ± | 0.17 | | | **0.42** | ± | 0.04 | | **0.43** | ± | 0.1 | | **0.44** | ± | 0.08 | **0.54** | | ± | 0.11 | **0.54** | | ± | 0.06 | **0.48** | | ± | 0.1 | **0.59** | | ± | 0.13 |
| Water level (cm) | CLT | **100.8** | | ± | 0.5 | | | **102.7** | ± | 0.3 | | **104.5** | ± | 0.7 | | **106.0** | ± | 1.3 | **106.8** | | ± | 2.0 | **108.4** | | ± | 0.6 | **107.5** | | ± | 4.4 | **109.2** | | ± | 0.3 |
|  | CHT | **100.5** | | ± | 0.7 | | | **101.2** | ± | 0.8 | | **101.8** | ± | 1.1 | | **102.2** | ± | 1.0 | **101.4** | | ± | 0.7 | **99.2** | | ± | 1.1 | **100.3** | | ± | 4.1 | **96.4** | | ± | 1.5 |
|  | TLT | **100.4** | | ± | 2.0 | | | **102.3** | ± | 1.3 | | **104.3** | ± | 1.0 | | **107.1** | ± | 2.0 | **107.4** | | ± | 1.2 | **108.0** | | ± | 0.7 | **109.1** | | ± | 0.2 | **109.3** | | ± | 0.5 |
|  | THT | **100.7** | | ± | 0.4 | | | **101.2** | ± | 0.6 | | **101.6** | ± | 1.0 | | **102.1** | ± | 1.2 | **101.4** | | ± | 0.7 | **99.6** | | ± | 1.1 | **98.6** | | ± | 1.4 | **96.1** | | ± | 1.6 |
| Chlorophyll-a (RFU) | CLT | **1.94** | | ± | 1.7 | | | **1.88** | ± | 1.96 | | **0.32** | ± | 0.17 | | **0.3** | ± | 0.1 | **0.30** | | ± | 0.1 | **0.34** | | ± | 0.25 | **0.45** | | ± | 0.42 | **0.09** | | ± | 0.11 |
|  | CHT | **0.73** | | ± | 0.9 | | | **0.66** | ± | 0.61 | | **0.38** | ± | 0.15 | | **0.35** | ± | 0.07 | **0.51** | | ± | 0.19 | **0.61** | | ± | 0.16 | **0.45** | | ± | 0.18 | **0.37** | | ± | 0.11 |
|  | TLT | **1.88** | | ± | 1.54 | | | **1.62** | ± | 1.48 | | **0.39** | ± | 0.29 | | **0.26** | ± | 0.08 | **0.32** | | ± | 0.12 | **0.50** | | ± | 0.25 | **0.55** | | ± | 0.33 | **0.11** | | ± | 0.09 |
|  | THT | **1.89** | | ± | 0.71 | | | **0.60** | ± | 0.19 | | **0.29** | ± | 0.08 | | **0.36** | ± | 0.06 | **0.53** | | ± | 0.16 | **0.83** | | ± | 0.23 | **0.48** | | ± | 0.12 | **0.34** | | ± | 0.09 |

**S6. Nutrient and phytoplankton community responses to warming and pharmaceuticals**

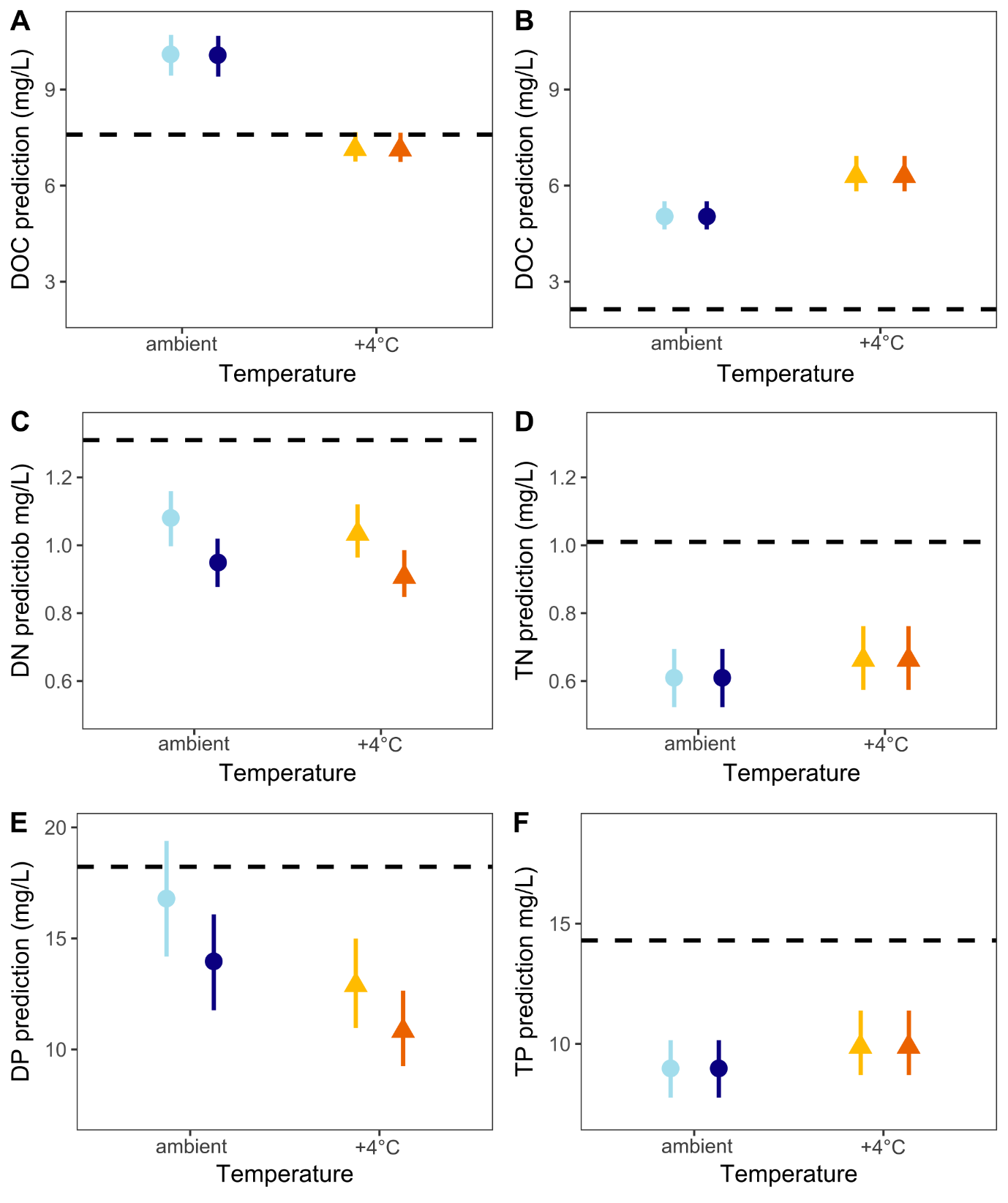

**Figure S6.1.** Most parsimonious models of treatment-specific concentrations of nutrients at the end of the summer (A, C, E) and winter (B, D, F) experiments. (A, B) dissolved organic carbon, DOC; (C) dissolved nitrogen, DN; (D) total nitrogen, TN; (E) dissolved phosphorus, SRP; (F) total phosphorus (TP). Black dashed line: initial values on Day 0. Treatments: light blue = non-heated, without pharmaceuticals; yellow = heated, without pharmaceuticals; dark blue = non-heated, with pharmaceuticals; orange = heated, with pharmaceuticals Model fits shown as mean values with 95% confidence intervals based on fixed effects.

**Table S6.1. Parameters of the most parsimonious Gamma GLM models of final nutrient concentrations in the summer experiment.** Values given on the predictor scale as fit with 95% confidence interval based on fixed effects only. Intercept = value for non-heated tanks without added pollutants; PhAC = difference in the mesocosms with added pharmaceuticals relative to untreated ones; Warming = difference in heated mesocosms relative to non-heated ones; ‘-‘ = effect not included in the model.

| **Parameter** | **DOC** | **DN** | **SRP** |
| --- | --- | --- | --- |
| (Intercept) | 2.31 (2.27–2.35) | 0.07 (0.03–0.11) | 2.80 (2.74–2.86) |
| PhAC | -0.01 (-0.05–0.03) | -0.13 (-0.17 to -0.09) | -0.18 (-0.25 to -0.11) |
| Warming | -0.33 (-0.38 to -0.29) | -0.03 (-0.07 – 0.01) | -0.24 (-0.32 to -0.16) |
| PhAC × Warming | - | - | - |
| $R_{c}^{2}$ / $R_{m}^{2}$ | - / 0.000 | - / 0.037 | - / 0.162 |

**Table S6.2. Parameters of the most parsimonious Gamma GLM models of final nutrient concentrations in the winter experiment.** Values and their interpretation as in Table S6.1.

| **Parameter** | **DOC** | **TN** | **TP** |
| --- | --- | --- | --- |
| (Intercept) | 1.62 (1.58–1.66) | -0.51 (-0.58 to -0.44) | 2.18 (2.12–2.24) |
| PhAC | - | - | -- |
| Warming | 0.23 (0.17–0.29) | 0.09 (-0.01– 0.19) | 0.11 (0.02–0.20) |
| PhAC × Warming | - | - | - |
| $R_{c}^{2}$ / $R_{m}^{2}$ | - / 0.000 | - / 0.037 | - / 0.162 |

Changes in the phytoplankton community (classes and lower-level taxa) over the experiment were tested by a redundancy analysis (RDA on centred data) separately for the summer and the winter experiment. Treatment was used as a categorical explanatory variable and community data (both class and species data) were used as a response. Statistical significance of the RDAs was tested by Monte Carlo permutation tests (999 unrestricted permutations under the reduced model). This analysis was run on data measured at the end of each experiment.

**Table S6.3. Results of the RDA analysis on the phytoplankton data sets.**

| Experiment | Group | Pseudo-F | *p* |
| --- | --- | --- | --- |
| Summer | Class | 0.6 | 0.89 |
|  | Taxon | 1.1 | 0.49 |
| Winter | Class | 1.4 | 0.19 |
|  | Taxon | 1.1 | 0.44 |

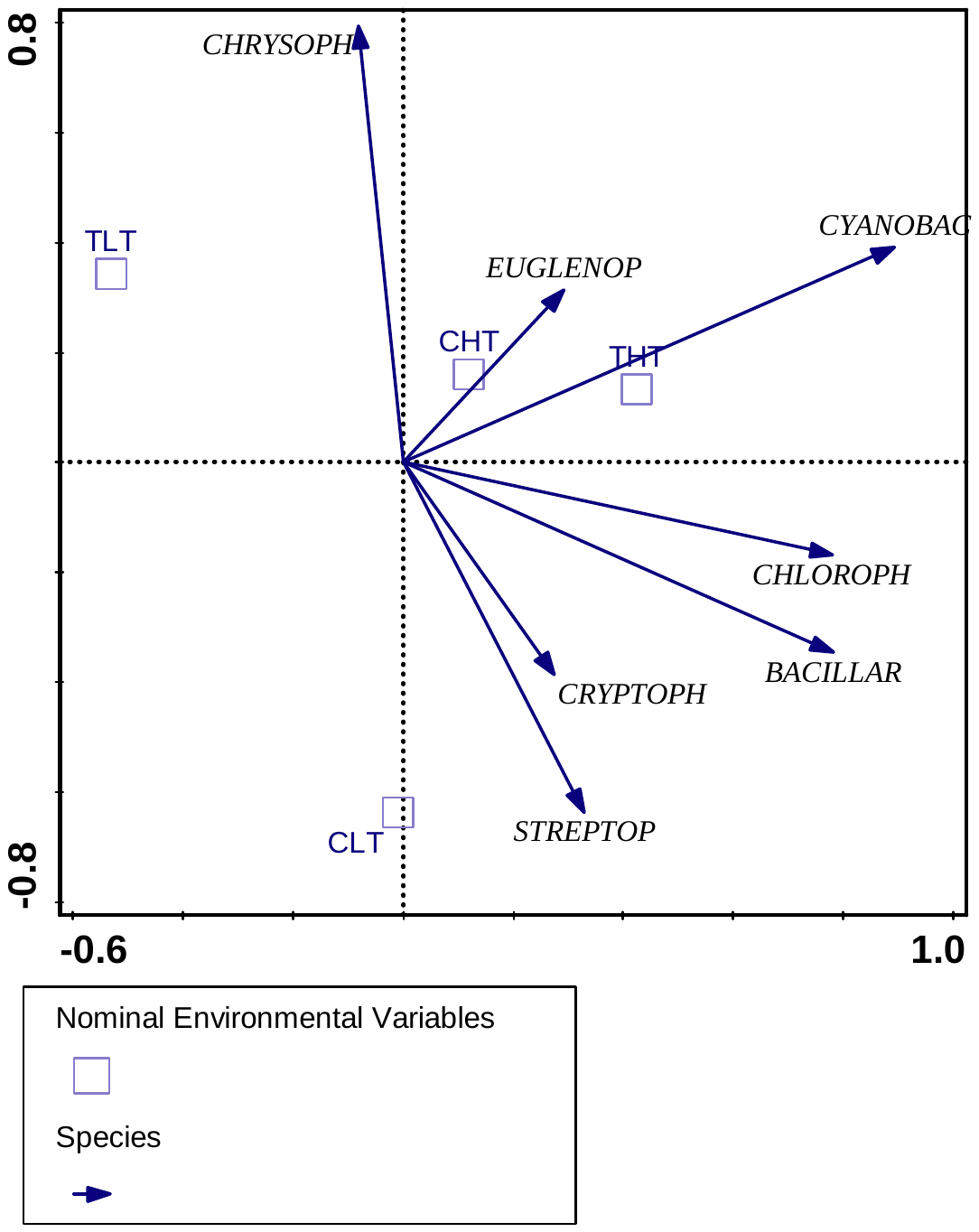

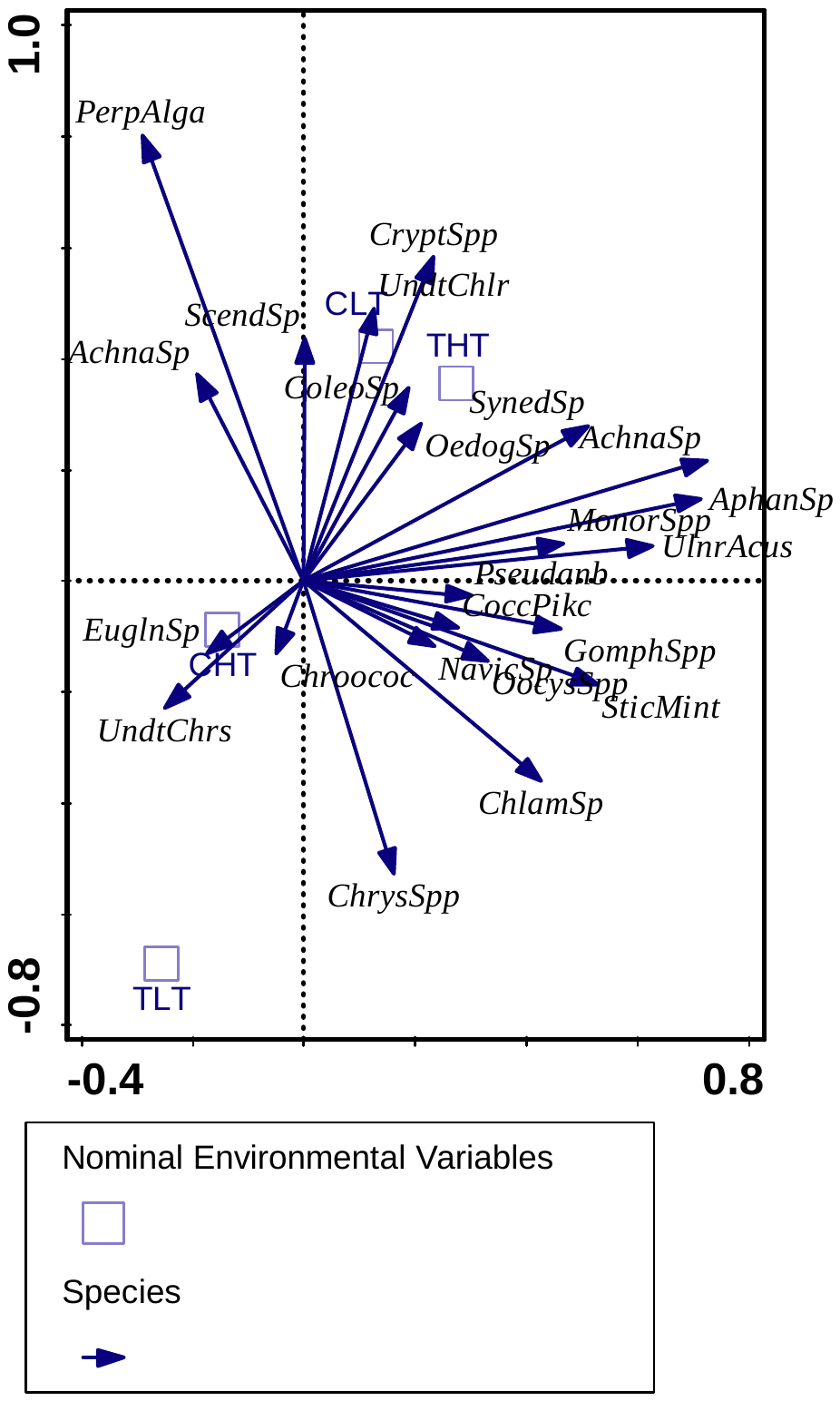

22.2 %

40.6 %

14.6 %

18.2 %

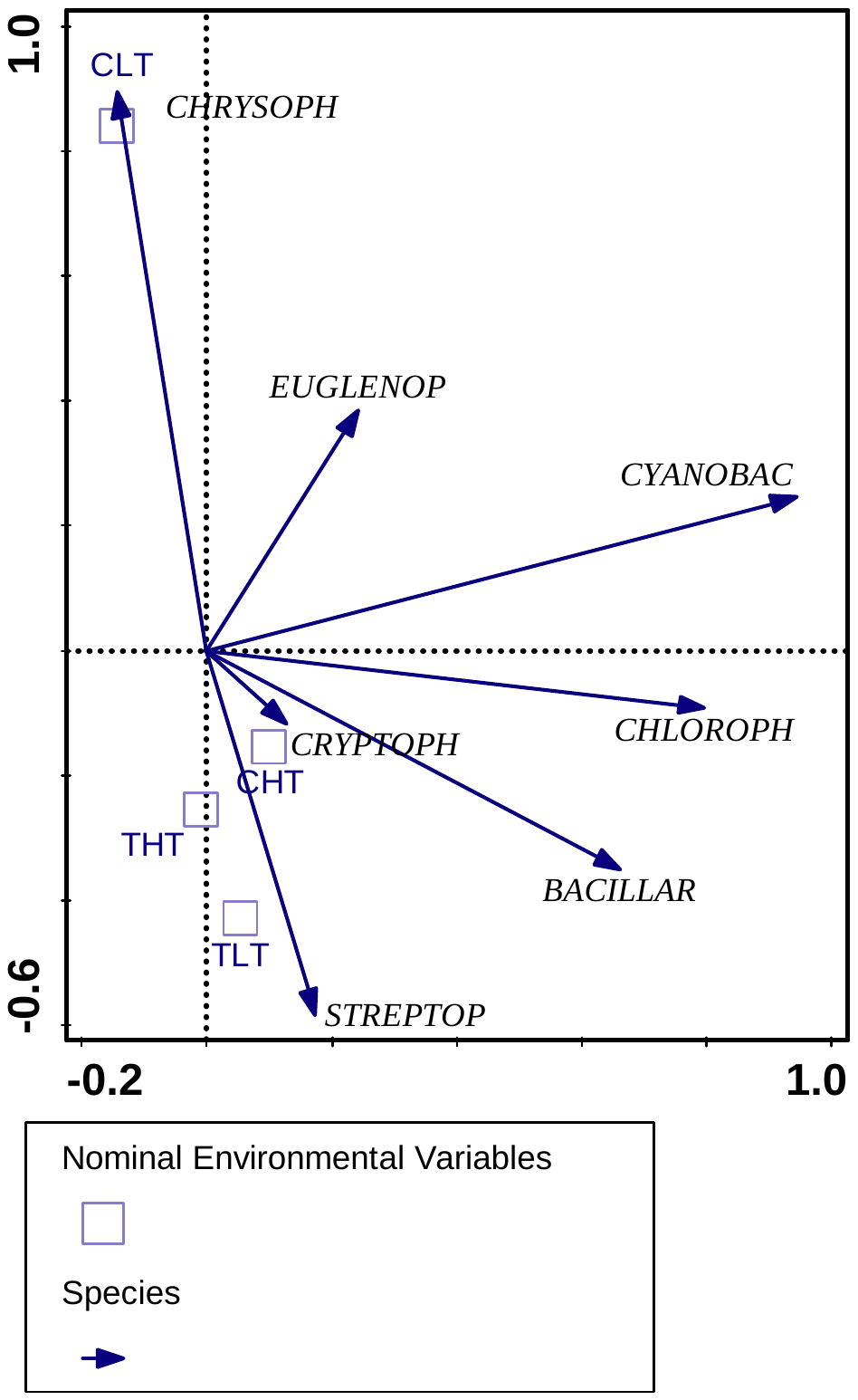

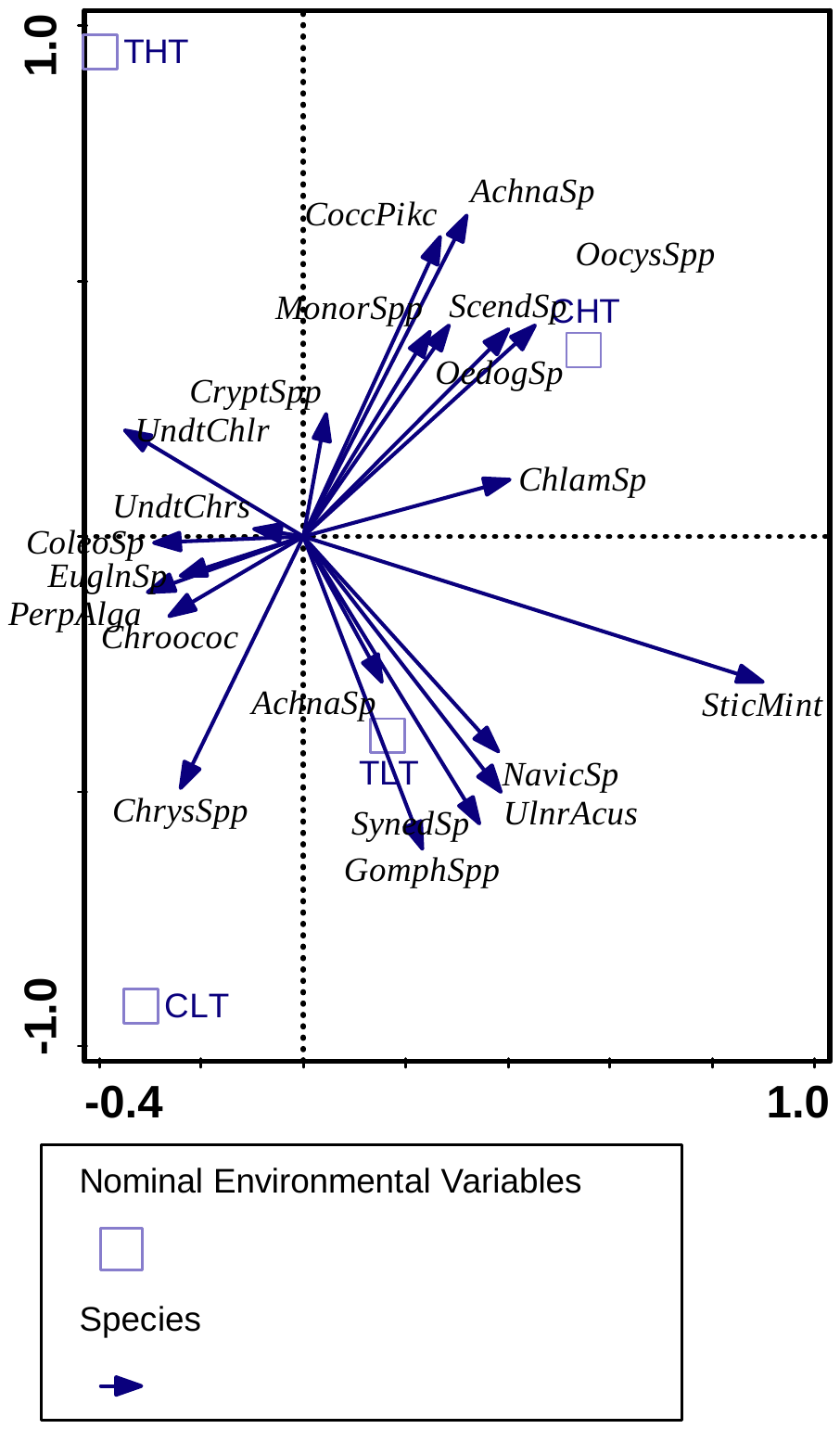

39.6 %

20.2 %

15.3 %

18.4 %

**Figure S6.2.** Unconstrained ordination diagrams (PCA) of algae main groups (A, C) and species community composition (B, D) in individual treatments in the summer (A, B) and winter (C, D) experiments. Explained variation (%) are represented next to each axis.

**S7. Zooplankton community responses to warming and pharmaceuticals**

**Table S7.1. Comparison of all candidate models of the temporal dynamics of *Daphnia* biomass during the summer and winter experiments.** Best model for each response variable in bold; ‘top model’ set with ΔAICc ≤ 2 for each response variable highlighted in bold italics. Time scaling = scaling used for the time variable (t). Model structure: *trt* = treatment effect, *t*, *t*^2^, *t*^3^ and *t*^4^ = time dependent effects; df = degrees of freedom; ln(*L*) = log-likelihood; ΔAICc = AICc difference from the best model; *w* = Akaike weight based on AICc; rank = model rank.

| Response | Time scaling | Model structure | df | ln(*L*) | ΔAICc | *w* | rank |
| --- | --- | --- | --- | --- | --- | --- | --- |
| *Daphnia* biomass (summer) | day | ***trt × t*** | **12** | **-393.6** | **0** | **0.082** | **1** |
|  | day | *trt ×* (*t+t^2^*) | 16 | -391.3 | 4.8 | 0.083 | 2 |
|  | day | *trt ×* (*t+t^2^+t^3^*) | 20 | -387.3 | 6.5 | 0.083 | 3 |
|  | day | *trt ×* (*t+t^2^+t3+t^4^*) | 24 | -385.3 | 12.8 | 0.083 | 8 |
|  | day ^0.5^ | *trt × t* | 12 | -412.8 | 38.6 | 0.086 | 12 |
|  | day ^0.5^ | *trt ×* (*t+t^2^*) | 16 | -393.5 | 9.1 | 0.083 | 6 |
|  | day ^0.5^ | *trt ×* (*t+t^2^+t^3^*) | 20 | -391.8 | 15.6 | 0.084 | 9 |
|  | day ^0.5^ | *trt ×* (*t+t^2^+t^3^+t^4^*) | 24 | -388.7 | 19.6 | 0.084 | 11 |
|  | day ^0.25^ | *trt × t* | 12 | -397.1 | 7.1 | 0.083 | 4 |
|  | day ^0.25^ | *trt ×* (*t+t^2^*) | 16 | -392.7 | 7.7 | 0.083 | 5 |
|  | day ^0.25^ | *trt ×* (*t+t^2^+t^3^*) | 20 | -390.3 | 12.6 | 0.083 | 7 |
|  | day ^0.25^ | *trt ×* (*t+t^2^+t^3^+t^4^*) | 24 | -386.8 | 15.8 | 0.083 | 10 |
| *Daphnia* biomass (winter) | day | *trt × t* | 12 | -355.7 | 187.4 | 0.096 | 10 |
|  | day | *trt ×* (*t+t^2^*) | 16 | -355.8 | 156.9 | 0.092 | 9 |
|  | day | *trt ×* (*t+t^2^+t^3^*) | 20 | -295.4 | 85.5 | 0.082 | 7 |
|  | day | *trt ×* (*t+t^2^+t^3^+t^4^*) | 24 | -261.8 | 28.1 | 0.074 | 2 |
|  | day ^0.5^ | *trt × t* | 12 | -355.9 | 187.8 | 0.096 | 11 |
|  | day ^0.5^ | *trt ×* (*t+t^2^*) | 16 | -300.7 | 86.7 | 0.082 | 8 |
|  | day ^0.5^ | *trt ×* (*t+t^2^+t^3^*) | 20 | -280.5 | 55.8 | 0.078 | 5 |
|  | day ^0.5^ | ***trt ×* (*t+t^2^+t^3^+t^4^*)** | **24** | **-247.7** | **0** | **0.071** | **1** |
|  | day ^0.25^ | *trt × t* | 12 | -361.2 | 198.5 | 0.097 | 12 |
|  | day ^0.25^ | *trt ×* (*t+t^2^*) | 16 | -288.0 | 61.1 | 0.079 | 6 |
|  | day ^0.25^ | *trt ×* (*t+t^2^+t^3^*) | 20 | -274.9 | 44.5 | 0.077 | 4 |
|  | day ^0.25^ | *trt ×* (*t+t^2^+t^3^+t^4^*) | 24 | -266.2 | 36.9 | 0.076 | 3 |

**Table S7.2. Comparison of all candidate models of the temporal dynamics of copepod biomass during the summer and winter experiments.** Best model for each response variable in bold; ‘top model’ set with ΔAICc ≤ 2 for each response variable highlighted in bold italics. Time scaling = scaling used for the time variable (*t*). Model structure: *trt* = treatment effect, *t*, *t*^2^, *t*^3^ and *t*^4^ = time-dependent effects; df = degrees of freedom; ln(*L*) = log-likelihood; ΔAICc = AICc difference from the best model; *w* = Akaike weight based on AICc; rank = model rank.

| Response | Time scaling | Model structure | df | ln(*L*) | ΔAICc | *w* | Rank |
| --- | --- | --- | --- | --- | --- | --- | --- |
| Copepod biomass (summer) | day | *trt × t* | 12 | -386.5 | 28.9 | 0.086 | 12 |
|  | day | ***trt ×* (*t+t^2^*)** | **16** | **-367.5** | **0** | **0.082** | **1** |
|  | day | *trt ×* (*t+t^2^+t^3^*) | 20 | -365.1 | 4.7 | 0.083 | 3 |
|  | day | *trt ×* (*t+t^2^+t^3^+t^4^*) | 24 | -359.5 | 3.5 | 0.082 | 2 |
|  | day ^0.5^ | *trt × t* | 12 | -376.1 | 8.0 | 0.083 | 7 |
|  | day ^0.5^ | *trt ×* (*t+t^2^*) | 16 | -372.8 | 10.6 | 0.084 | 9 |
|  | day ^0.5^ | *trt ×* (*t+t^2^+t^3^*) | 20 | -367.0 | 8.6 | 0.083 | 8 |
|  | day ^0.5^ | *trt ×* (*t+t^2^+t^3^+t^4^*) | 24 | -364.3 | 13.1 | 0.084 | 11 |
|  | day ^0.25^ | *trt × t* | 12 | -375.3 | 6.4 | 0.083 | 6 |
|  | day ^0.25^ | *trt ×* (*t+t^2^*) | 16 | -370.2 | 5.5 | 0.083 | 4 |
|  | day ^0.25^ | *trt ×* (*t+t^2^+t^3^*) | 20 | -365.6 | 5.7 | 0.083 | 5 |
|  | day ^0.25^ | *trt ×* (*t+t^2^+t^3^+t^4^*) | 24 | -363.6 | 11.7 | 0.083 | 10 |
| Copepod biomass (winter) | day | *trt × t* | 12 | -149.9 | 39.8 | 0.086 | 6 |
|  | day | *trt ×* (*t+t^2^*) | 16 | -148.6 | 46.2 | 0.087 | 9 |
|  | day | *trt ×* (*t+t^2^+t^3^*) | 20 | -129.0 | 16.5 | 0.079 | 4 |
|  | day | *trt ×* (*t+t^2^+t^3^+t^4^*) | 24 | -120.1 | 8.6 | 0.076 | 2 |
|  | day ^0.5^ | *trt × t* | 12 | -158.9 | 57.8 | 0.091 | 12 |
|  | day ^0.5^ | *trt ×* (*t+t^2^*) | 16 | -146.7 | 42.5 | 0.086 | 7 |
|  | day ^0.5^ | *trt ×* (*t+t^2^+t^3^*) | 20 | -146.0 | 50.5 | 0.088 | 11 |
|  | day ^0.5^ | ***trt ×* (*t+t^2^+t^3^+t^4^*)** | **24** | **-115.8** | **0** | **0.074** | **1** |
|  | day ^0.25^ | *trt × t* | 12 | -153.6 | 47.2 | 0.088 | 10 |
|  | day ^0.25^ | *trt ×* (*t+t^2^*) | 16 | -147.9 | 44.8 | 0.087 | 8 |
|  | day ^0.25^ | *trt ×* (*t+t^2^+t^3^*) | 20 | -133.1 | 24.7 | 0.081 | 5 |
|  | day ^0.25^ | *trt ×* (*t+t^2^+t^3^+t^4^*) | 24 | -121.5 | 11.4 | 0.077 | 3 |

**Table S7.3. Parameters of the best GLMM models describing the temporal dynamics of Daphnia and copepod biomass characteristics in the summer experiment.** Model structure: zero-inflated Gamma with log-link function for *Daphnia* biomass and Gamma with log-link function for copepod biomass. Fixed effect parameter estimates given as mean with 95% confidence interval in parentheses. Intercept corresponds to day 0 in controls; “PhAC” (pharmaceuticals) and “Warming” describe differences of the given treatment from controls. Random effects: *σ*^2^ = residual variance; *τ*_00_ = random effect variance; *R^2^_m_* = marginal *R^2^*, *R^2^_c_* = conditional *R^2^*; all models with *N* = 28 random effect levels (mesocosms) and *N_tot_* = 196 observations.

| **Parameter** | ***Daphnia* biomass** | **Copepod biomass** |
| --- | --- | --- |
| (Intercept) | 1.60 (1.40–1.80) | 1.06 (0.96–1.16) |
| PhAC | -0.35 (-0.58 to -0.13) | - |
| Warming | -0.88 (-1.11 to -0.66) | - |
| *day* | -5.51 (-8.56 to -2.46) | 8.23 (5.66–10.79) |
| *day*^2^ | - | -5.39 (-7.87 to -2.91) |
| PhAC × *day* | -9.04 (-12.05 to -6.02) | 8.35 (5.73–10.96) |
| PhAC × *day*^2^ | - | -4.97 (-7.45 to -2.48) |
| Warming × *day* | -7.82 (-11.18 to -4.45) | - 6.98 (-10.46 to -3.50) |
| Warming × *day*^2^ | - | 4.69 (1.35–8.03) |
| PhAC × Warming × *day* | - | 6.37 (1.35–11.39) |
| PhAC × Warming × *day*^2^ | - | -2.59 (-7.34–2.16) |
| **Zero-Inflated Model** | | |
| (Intercept) | -5.70 (-7.59 to -3.80) | - |
| Day | 0.10 (0.06–0.14) | - |
| **Random Effects** | | |
| σ^2^ | 0.53 | 0.40 |
| τ_00_ | 0.00 _mesocosm_ | 0.02 _mesocosm_ |
| *R^2^_m_* / *R^2^_c_* | 0.646 / NA | 0.447 / 0.468 |

**Table S7.4. Parameters of the best GLMM models describing the temporal dynamics of *Daphnia* and copepod biomasses in the winter experiment.** Model structure: zero-inflated Gamma with log-link function for *Daphnia* and copepod biomass. Fixed effect parameter estimates given as mean with 95% confidence interval in parentheses. Intercept corresponds to day 0 in controls; “PhAC” (pharmaceuticals) and “Warming” describe differences of the given treatment from controls. Random effects: σ2 = residual variance; τ00 = random effect variance; *R^2^_m_* = marginal *R^2^*, *R^2^_c_* = conditional *R^2^*; all models with *N* = 28 random effect levels (mesocosms) and *N_tot_* = 224 observations.

| **Parameter** | ***Daphnia* biomass** | **Copepod biomass** |
| --- | --- | --- |
| (Intercept) | 0.44 (0.35–0.52) | -0.44 (-0.57 – -0.32) |
| *day*^0.25^ | 5.07 (3.65–6.48) | -1.43 (-3.16–0.29) |
| *day*^0.5^ | -6.69 (-8.13 to -5.25) | -2.53 (-4.26 to -0.80) |
| *day*^0.75^ | -6.25 (-7.69 to -4.80) | -0.87 (-2.64–0.91) |
| *day* | 5.46 (4.03–6.89) | 5.64 (3.87–7.42) |
| Warming × *day*^0.25^ | 2.69 (1.30–4.07) | - |
| Warming × *day*^0.5^ | -6.75 (-8.12 to -5.38) | - |
| Warming × *day*^0.75^ | -2.39 (-3.76 to -1.03) | - |
| Warming × *day* | 2.79 (1.41–4.16) | - |
| PhAC × *day*^0.25^ | - | -3.55 (-5.23 to -1.87) |
| PhAC × *day*^0.5^ | - | -3.39 (-5.06 to -1.71) |
| PhAC × *day*^0.75^ | - | -1.29 (-2.92–0.35) |
| PhAC × *day* | - | 4.70 (2.99–6.41) |
| **Zero-Inflated Model** |  |  |
| (Intercept) | -4.01 (-5.68 to -2.33) | -3.33 (-4.36 to -2.31) |
| *Day* | -0.019 (-0.063–0.025) | 0.001 (-0.011–0.012) |
| **Random Effects** |  |  |
| σ^2^ | 0.25 | 0.35 |
| τ_00_ | 0.03 _mesocosm_ | 0.06 _mesocosm_ |
| *R^2^_m_* / *R^2^_c_* | 0.627 / 0.662 | 0.325 / 0.423 |

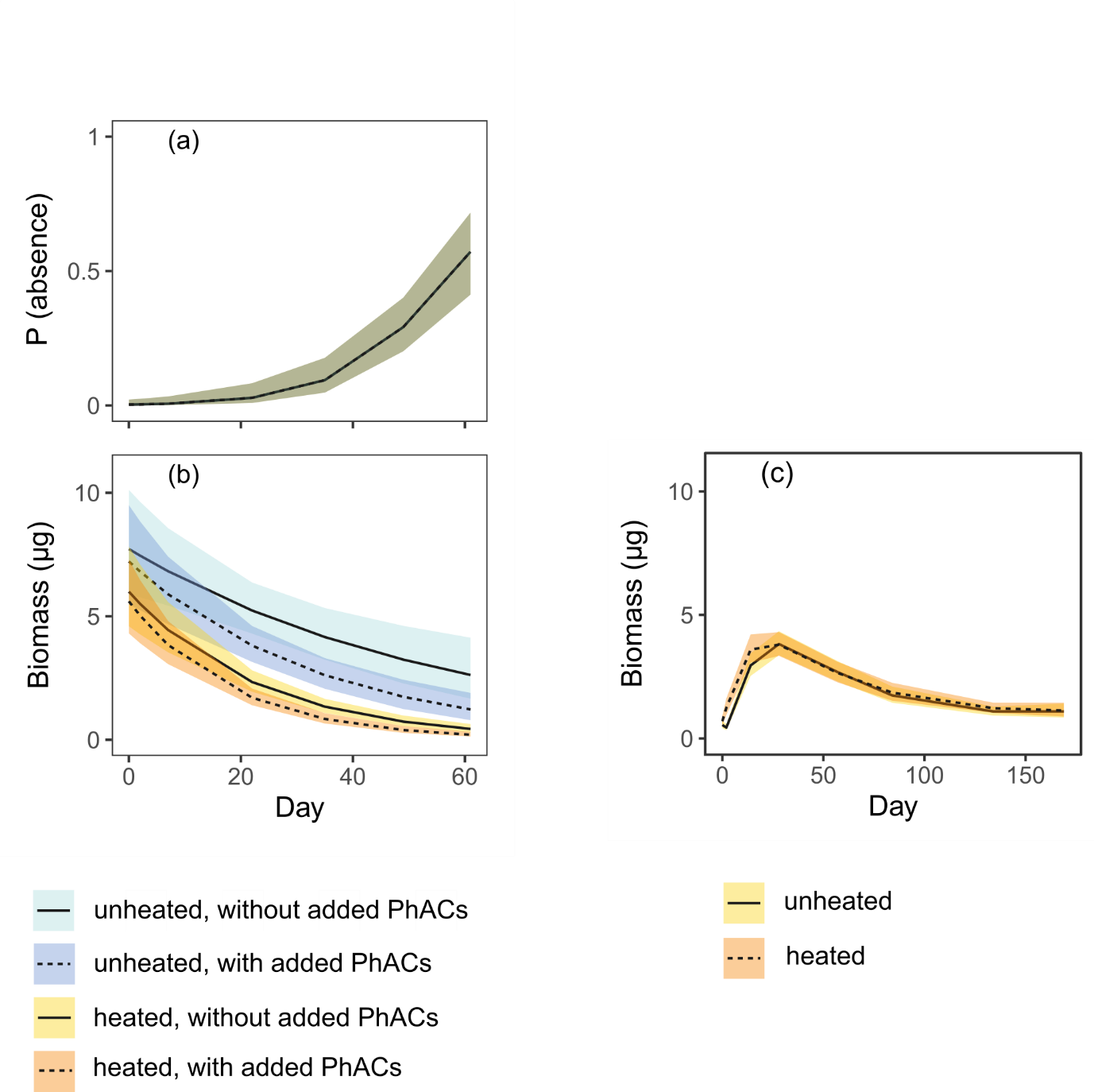

**Figure S7.1.** Most parsimonious models of treatment-specific *Daphnia* biomass during the summer (a, b) and winter (c) experiments. Model estimates shown as mean values with 95% confidence intervals based on fixed effects. For the data collected in summer, predictions for biomass are separated into (a) the probability of *Daphnia* absence based on the zero-inflated Gamma generalized linear mixed effect model (GLMM) and (b) *Daphnia* biomass based on the conditional Gamma GLMM. *Daphnia* were absent in only two samples in winter and the probability of their absence based on the zero-inflated Gamma GLMM is not shown. Treatment: PhACs = pharmaceuticals.

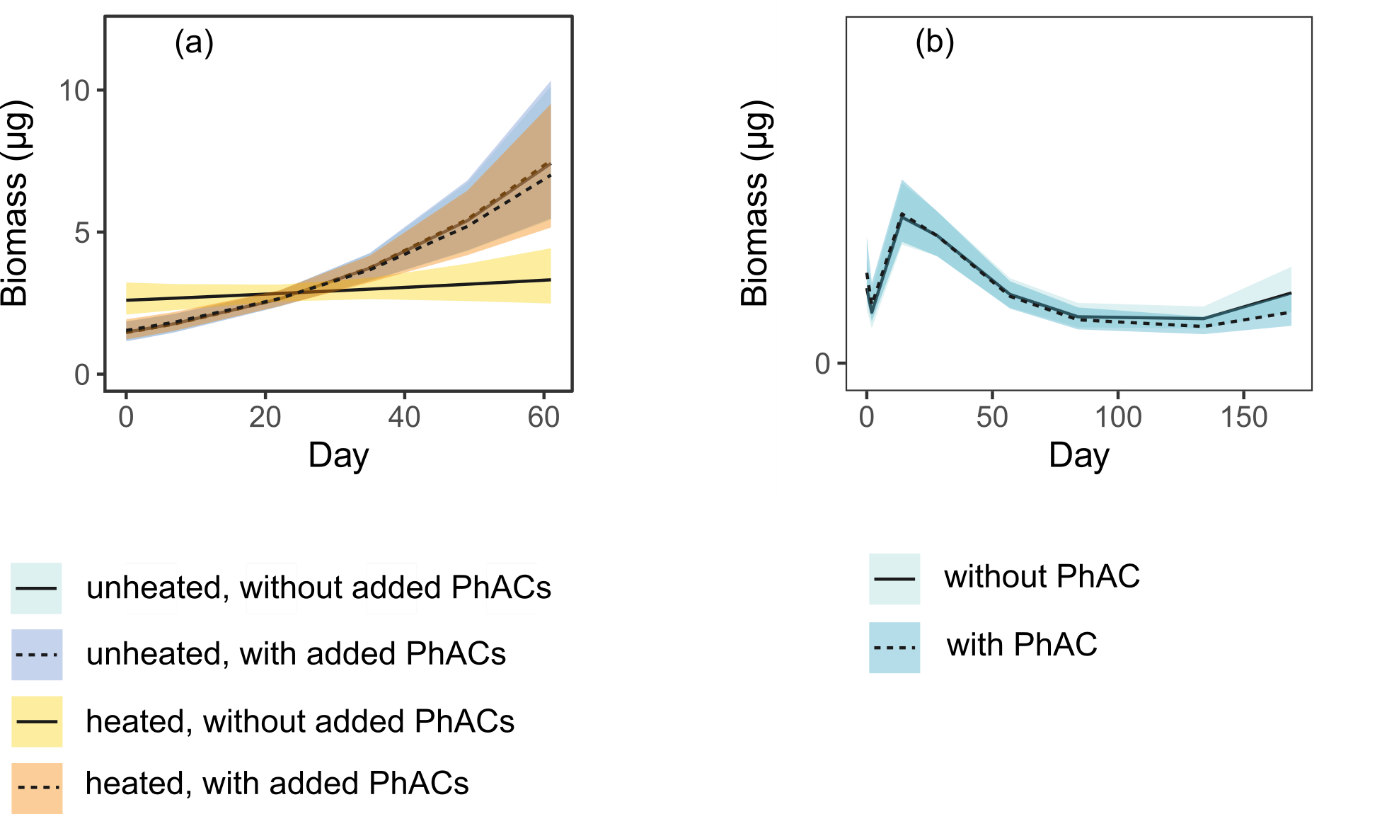

**Figure S7.2**. Most parsimonious models of treatment-specific copepod biomass during the summer (a) and winter (b) experiments. Model estimates shown as mean values with 95% confidence intervals based on fixed effects. In summer and winter, copepods were absent in only four samples and the probability of their absence based on the zero-inflated Gamma GLMM is not shown. Treatment: PhACs = pharmaceuticals.

**S8. Insect predator responses to warming and pharmaceuticals**

**Table S8.1. Parameters of the most parsimonious binomial GLM models of cumulative odonate emergence in the summer experiment.** Values and their interpretation as in Table S6.1.

| **Parameter** | ***Aeshna*** | **Zygoptera** |
| --- | --- | --- |
| (Intercept) | 1.22 (0.70–2.12) | 0.72 (0.55–0.94) |
| PhAC | 3.00 (1.44–6.26) | - |
| Warming | 2.05 (1.00–4.21) | 0.60 (0.41–0.89) |
| PhAC × Warming | - | - |
| $R_{c}^{2}$ / $R_{m}^{2}$ | - / 0.119 | - / 0.020 |

**Table S8.2. Parameters of the most parsimonious binomial GLM models of the survival probability of zygopteran and *Anax* larvae and adult *Notonecta* in the winter experiment.** Values and their interpretation as in Table S6.1.

| **Parameter** | **Zygoptera** | ***Notonecta*** | ***Anax*** |
| --- | --- | --- | --- |
| (Intercept) | 0.96 (0.74–1.25) | 1.05 (0.68–1.61) | 0.87 (0.64–1.17) |
| PhAC | 1.69 (1.16–2.47) | - | - |
| Warming | 2.06 (1.41–3.02) | 0.22 (0.11–0.44) | - |
| PhAC × Warming | 0.26 (0.15–0.44) | - | - |
| $R_{c}^{2}$ / $R_{m}^{2}$ | - / 0.037 | - / 0.150 | NA / 0.000 |

**S9. Effects of the stressors on the food web**

**Table S9.1**. **Estimates of regression coefficients, covariances and variables for the structural equation model of the relationships between stressors and community components in the summer experiment.** Treatment: PhAC = pharmaceuticals. Significant predictors in bold. Model diagnostics: Fisher's *C* = 2.55, *p* = 0.28, df = 2.

| Response | Predictor | Estimate | SE | *P* |
| --- | --- | --- | --- | --- |
| Filter feeders | **PhAC** | **< -0.01** | **0.00** | **0.047** |
|  | **Warming** | **-0.14** | **0.02** | **< 0.001** |
|  | Phytoplankton | 0.09 | 0.22 | 0.669 |
| Phytoplankton | **PhAC** | **< -0.01** | **0.00** | **0.032** |
|  | **Warming** | **0.08** | **0.00** | **< 0.001** |
| Predators | PhAC | < 0.01 | 0.00 | 0.10 |
|  | **Warming** | **0.02** | **0.08** | **< 0.001** |
|  | **Filter feeders** | **-0.21** | **0.11** | **< 0.001** |

**Table S9.2**. **Estimates of regression coefficients, and variables for the structural equation model of the relationships between stressors and community components in the winter experiment.** Treatment: PhAC = pharmaceuticals. Significant predictors in bold. Model diagnostics: Fisher's *C* = 5.20, *p* = 0.074, df = 2.

| Response | Predictor | Estimate | SE | *P* |
| --- | --- | --- | --- | --- |
| Filter feeders | PhAC | < 0.01 | 0.00 | 0.127 |
|  | Warming | 0.06 | 0.09 | 0.478 |
|  | Phytoplankton | -1.25 | 1.22 | 0.305 |
| Phytoplankton | PhAC | < 0.01 | 0.00 | 0.798 |
|  | **Warming** | **0.28** | **0.09** | **0.002** |
| Predators | PhAC | < -0.01 | 0.00 | 0.525 |
|  | Warming | < -0.01 | 0.02 | 0.935 |
|  | Filter feeders | 0.09 | 0.08 | 0.291 |

Ter Braak, C.J.F., Smilauer, P., 2012. Canoco reference manual and user’s guide: software for ordination, version 5.0.

Van den Brink, P.J., ter Braak, C.J.F., 1999. Principal response curves: Analysis of time-dependent multivariate responses of biological community to stress. Environ. Toxicol. Chem. 18, 138–148. https://doi.org/10.1002/etc.5620180207

Widbom, B. 1984. Determination of average individual dry weights and ash-free dry weights in different sieve fractions of marine meiofauna. Marine Biology 84: 101-108.

Yıldız D., Yalçın G., Jovanović B., Boukal B.S., Vebrová L., Riha D., Stanković J., Savić-Zdraković D., Metin M., Akyürek Y. N., Balkanlı D., Filiz N., Milošević D., Feuchtmayr H., Richardson J.A., Beklioğlu M. (2022) Effects of a microplastic mixture differ across trophic levels and taxa in a freshwater food web: In situ mesocosm experiment. Science of The Total Environment. <https://doi.org/10.1016/j.scitotenv.2022.155407>.
